# Optimizing therapeutic hypothermia conditions in a translational preclinical model of neonatal hypoxia-ischemia in rats

**DOI:** 10.64898/2026.02.06.704040

**Authors:** Ifrah Omar Ibrahim, Pierre Goudeneche, Léonie Dayraut, Stéphane Sanchez, Jean Delmas, Jean-François Chateil, Luc Pellerin, Hélène Roumes, Anne-Karine Bouzier-Sore

**Author notes:** Corresponding author : Anne-Karine Bouzier-Sore, Univ. Bordeaux, CNRS, CRMSB, UMR 5536, F-33000 Bordeaux, France. These authors contributed equally to this work (Ifrah Omar Ibrahim, Pierre Goudeneche). These authors also contributed equally to this work (Hélène Roumes, Anne-Karine Bouzier-Sore).

## Abstract

**Background:** Therapeutic hypothermia is the only clinically approved treatment for neonatal hypoxia–ischemia (NHI), although its efficacy remains partial. In preclinical research, hypothermia is widely used as a reference therapy; however, its protocol is highly variable across studies, limiting robust comparisons with emerging neuroprotective strategies. This study aimed to define an optimal and standardized hypothermia protocol in the Rice–Vannucci model, not to challenge clinical practice, but to establish a reliable benchmark for preclinical therapeutic development.

**Methods:** NHI was induced in postnatal day 7 (P7) rat pups, followed by normothermia or hypothermia for 2, 3, or 5 hours. Short- and long-term outcomes were assessed using lesion volume measurements by MRI, neurological scoring, behavioral tests, and histological analyses. The impact of immediate hypothermia initiation was also examined.

**Results:** Across analyses, both 2- and 3-hour hypothermia durations provided greater neuroprotection than 5 hours—including brain lesion volume, motor and cognitive performances, and markers of neuronal preservation and neuroinflammation. However, for several parameters, 2 hours of hypothermia showed superior efficacy compared with 3 hours. Immediate initiation further modestly improved outcomes.

**Conclusion:** A 2-hour hypothermia protocol represents the most robust and reproducible preclinical reference, enabling meaningful comparison with novel therapies in the Rice–Vannucci model.

**IMPACT:**

- By establishing an optimized hypothermia protocol in the Rice–Vannucci model, this study offers a consistent and robust reference for preclinical evaluation of emerging therapies.
- It does not question clinical hypothermia protocols, but addresses variability in preclinical literature
- Optimizing the hypothermia reference protocol is mandatory to reliably identify new effective treatments in preclinical studies and to enhance their likelihood of successful and efficient clinical translation

## INTRODUCTION

The perinatal period represents a phase of heightened vulnerability. After prematurity, neonatal hypoxia-ischemia (NHI) is the main causes of perinatal mortality, affecting 2 to 6‰ of births in developed nations and up to 26‰ in underdeveloped countries ^1,2^. Therefore, NHI accounts for around 600,000 neonatal deaths each year, with at least as many infants experiencing severe motor and cognitive impairments ^3^. NHI is marked by a significant reduction in cerebral blood flow and a failure to deliver oxygen and energy sources (such as glucose and ketone bodies). This event is followed by two periods of energetic deficit separated by a latency phase, which corresponds to the therapeutic window ^4^. The only neuroprotective intervention that can improve outcomes for neonates after an hypoxic-ischemic event is therapeutic hypothermia ^5^. However, its application is limited to moderate to severe hypoxic-ischemic episodes in neonates older than 36 weeks of gestational age, due to a high risk of collateral damages such as intracerebral hemorrhages before this term ^6^. It should also be noted that this therapy is effective only after acute perinatal events, such as placental abruption or umbilical cord prolapse, and does not improve the prognostic if the hypoxic-ischemic insult occurred during the antenatal period or is due to prenatal chronic insults ^7^. Therapeutic hypothermia can be performed either through whole-body cooling or selective head cooling (33 ± 1°C, for 72 hours, followed by slow and gradual rewarming of the infant over 6 to 12 hours ^8,9^). Both methods are used in clinical settings without preference, as neither has demonstrated superior outcomes ^10,11^. While whole-body cooling may be more effective for achieving deep brain cooling ^12^, selective head cooling can limit the adverse effects of systemic cooling ^13^, even though these effects are reversible and generally moderate compared to the benefits. Common side effects of hypothermia include sinus bradycardia (a decrease in heart rate of 14 beats per minute is observed per degree below 37°C), arterial hypotension due to hypovolemia (which may require the administration of inotropes), mild thrombocytopenia, and pulmonary hypertension ^14^. Although the side effects of therapeutic hypothermia are reversible, this treatment has limitations in eligible neonates, as 44-53% do not respond favorably ^15^. Therefore, it is necessary to develop safe, simple, and effective adjunct therapies to complement the current treatment strategies for infants with NHI. The mechanisms involved in the neuroprotective effects of hypothermia are multifactorial. In a preclinical model of NHI in lambs, it was first demonstrated that neuroprotection of therapeutic hypothermia acts by preventing secondary cytotoxic edema and reducing neuronal loss ^16^. Lowering the baby’s temperature reduces brain energy metabolism by 5% to 8% for each degree of temperature decrease ^17^. This reduction in oxidative metabolism has been associated with a reduction of free radical production, as well as a decrease in membrane lipid peroxidation ^18^. Furthermore, during the latency phase (around 6 hours post-hypoxia-ischemia), therapeutic hypothermia has been shown to decrease pro-apoptotic caspase-3 activity, thereby limiting neuronal death ^19^. Reducing brain temperature also helps mitigate neuroinflammatory reactions, thus protecting mitochondrial oxidative phosphorylation ^20^. Numerous experimental studies have since demonstrated the neuroprotective role of hypothermia in the context of NHI. In P7 (7 days post-natal) rat pups exposed to a hypoxic-ischemic event, moderate hypothermia (31 to 34°C) significantly reduced cortical brain damage ^21^, as well as injury to the hippocampus, basal ganglia and thalamus ^22^, which correlated with improved sensorimotor recovery ^23^. Comparable outcomes have been reported in lambs ^24^ and piglets ^25^, species that are more neurodevelopmentally similar to humans ^26,27^. However, while most preclinical data suggest a neuroprotective role for hypothermia, a decrease in brain temperature (−2°C) could induce physiological stress, leading to prolonged elevation of circulating cortisol levels and, ultimately, increased neuronal loss, as demonstrated in lambs ^28,29^.

There is significant variability in the conditions under which hypothermia is implemented across preclinical studies. Specifically, focusing on studies involving rats, the most commonly used animal model for evaluating therapeutic strategies, the primary durations for hypothermia in the context of NHI are 2 hours ^30^, 3 hours ^21,22,31–33^, or 5 hours ^34–39^. Regrettably, the different durations were evaluated across separate studies, each conducted under distinct experimental conditions. This heterogeneity makes it difficult to directly compare the results or to establish the optimal duration for therapeutic hypothermia. As a result, the optimal duration for achieving maximal neuroprotection remains uncertain, complicating the selection of appropriate parameters for the efficient evaluation and comparison of our experimental treatment. Notably, one study in the Rice–Vannucci Sprague Dawley rat model performed a longitudinal comparison using MRI to assess 24 h versus 48 h of hypothermia ^40^, highlighting the value of imaging for temporal monitoring. However, shorter durations of 2, 3, or 5 hours remain the most commonly employed in preclinical research, which is why the present study systematically compares these durations under consistent conditions to determine the hypothermia protocol that provides maximal neuroprotection. Furthermore, in line with the new standards of animal experimentation, which emphasize compliance with the 3Rs (Replacement, Reduction, and Refinement), our study aims to define the threshold of hypothermia that allows for the measurement of beneficial effects, while minimizing potential pain, suffering, or distress, thereby enhancing animal welfare. Lastly, this work adheres to point No. 4 of the STAIR consortium recommendations (Stroke Therapy Academic Industry Roundtable): “*Outcome measures should include both infarct volume and functional assessment in both acute and long-term phase animal studies*” ^41^. Accordingly, the effect of therapeutic hypothermia was evaluated from the neonatal stage (P7) to adulthood (P51), at the anatomical level by non-invasive measurement of brain hypoxic-ischemic lesion volumes, using Magnetic Resonance Imaging (MRI), and at the functional level through behavioral assessments, including early reflexes, modified Neurological Severity Score, novel object recognition, anxiety and depression. Finally, histological and immunohistochemical analyses were performed to complement the imaging and behavioral data.

## MATERIALS AND METHODS

### Study type

This study is a pre-clinical study. The experimental design and groups are presented in Fig. 1. For this study, 6 groups of rat pups were considered, based on the hypoxia-ischemia event and the duration of therapeutic hypothermia. In the Sham group, the left common carotid artery was only exposed, without a hypoxia-ischemia insult or the application of hypothermia, and the animals were maintained at normothermia for 2 hours after MRI study. The control group (HIN group) underwent the hypoxic-ischemic event and MRI examinations, while remaining at normothermia for 2 hours post-MRI. Three groups underwent the hypoxic-ischemic event as well as MRI examinations and were subsequently subjects to therapeutic hypothermia for 2 hours, 3 hours, and 5 hours, respectively (HIH2, HIH3, and HIH5 groups). The last group underwent hypoxia-ischemia and was immediately subjected to hypothermia for 2 hours, after leaving the hypoxic chamber (without MRI at P7) (HIH2Im group).

**Figure 1:**
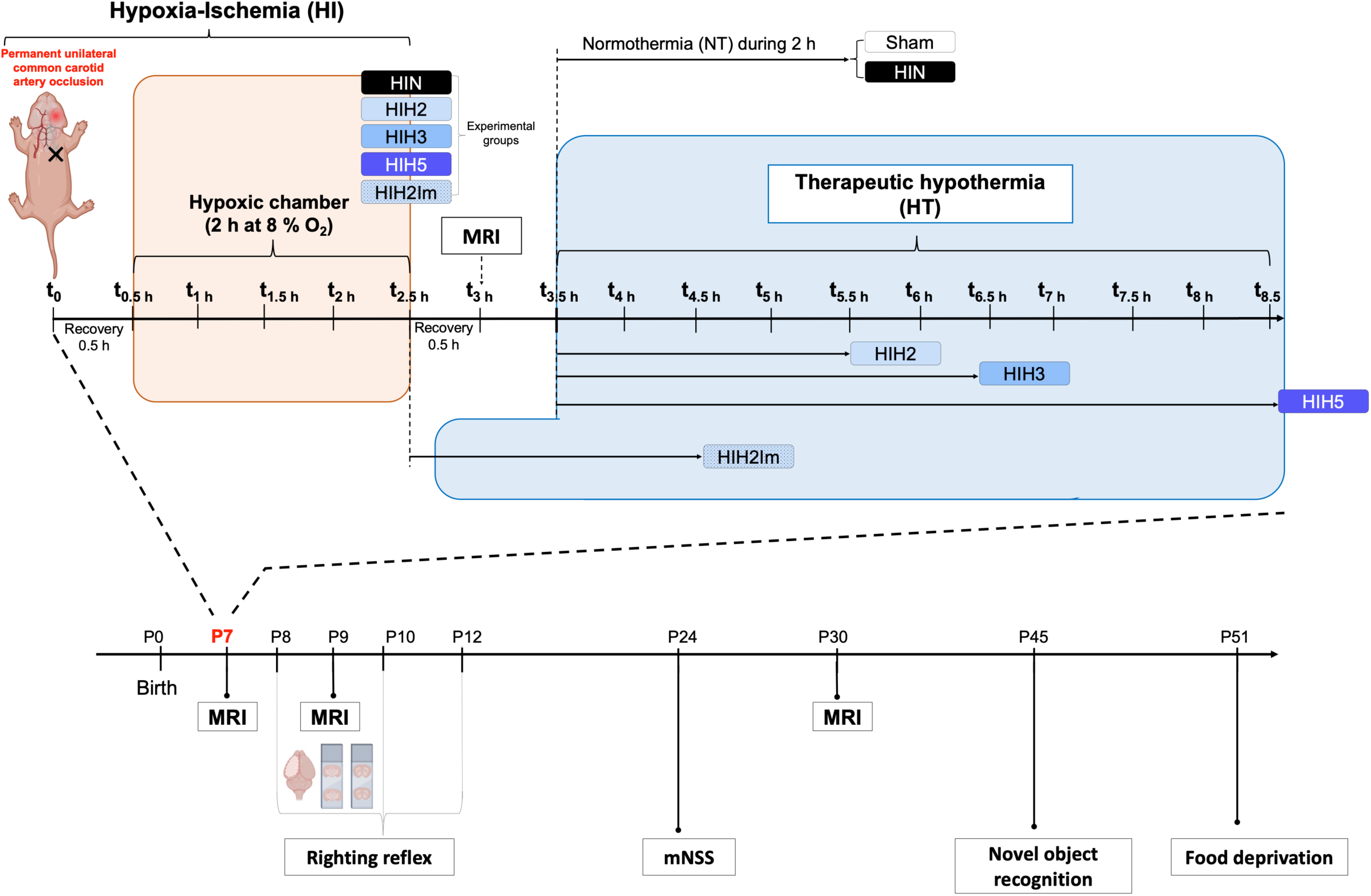
Experimental design and groups. Postnatal day 7 (P7) Wistar rat pups (n = 67) were randomly assigned to six experimental groups. Sham group (n = 13) underwent exposure of the left common carotid artery without ligation, with no hypoxia exposure, followed by 2 hours of controlled normothermia. Hypoxic-ischemic normothermia group (HIN, n = 14) underwent ligation of the left common carotid artery followed by 2 hours of hypoxia (8% O₂), magnetic resonance imaging (MRI) 3 hours post-surgery, and then 2 hours of controlled normothermia. Hypothermia 2-hour treatment group (HIH2, n = 11) underwent the hypoxic-ischemic protocol as above, MRI 3 hours post-surgery, followed by 2 hours of controlled therapeutic hypothermia. Hypothermia 3-hour treatment group (HIH3, n = 11): same hypoxic-ischemic protocol and MRI, followed by 3 hours of therapeutic hypothermia. Hypothermia 5-hour treatment group (HIH5, n = 11): same hypoxic-ischemic protocol and MRI, followed by 5 hours of therapeutic hypothermia. Immediate hypothermia group (HIH2Im, n = 7) underwent the hypoxic-ischemic protocol immediately followed by 2 hours of controlled therapeutic hypothermia (no delay and no MRI before cooling). The longitudinal evolution of brain lesion volume was assessed by MRI at P9 and P30. Behavioral testing included the righting reflex at P8, P10, and P12; the modified Neurological Severity Score (mNSS) at P24; and cognitive assessments at P45 for memory performance and at P51 for anxiety and depressive-like behaviors.

### Animals

This study was carried out in strict accordance with the recommendations in the Guide for the Care and Use of Laboratory Animals of the European Communities Council Directive of the 24 November 1986 (86/609/EEC). Protocols were consistent with the ethical guidelines of the French Ministry of Agriculture and Forest and the ARRIVE guidelines and received approval by the Committee on the Ethics of Animal Experiments of the University of Bordeaux (Protocol Number C2EA-50, authorization n°37955). Pregnant Wistar RJ-HAN females (Charles River, France) were received on gestational day 16 and were housed in standard environmental conditions with a 12:12h light:dark cycle and access to food and water *ad libitum*. They were housed in ventilated polycarbonates cages with sawdust bedding, at controlled room temperature (21-22°C) and humidity (55-60%). The animals were randomly assigned to an experimental group. Within the same litter (maximum of 8 pups per litter), the durations of hypothermia were randomized. Randomization was performed using a stratified design to evenly distribute treatment conditions across litters and to minimize litter effects. All efforts were made to minimize animal suffering. Surgical procedures and MRI exams were performed under isoflurane anesthesia with careful monitoring of physiological parameters including breath and body temperature. After surgery, pups were monitored daily for general health and welfare indicators, including weight gain, grooming behavior, spontaneous activity, and sign of pain or distress. In neonatal pups, lack of weight gain over 48 hours was considered a critical welfare threshold, along with other endpoints such as lethargy, abnormal posture, or dehydration. At the end of the experimental protocol, animals were first anesthetized with isoflurane (4%), and then received an intraperitoneal injection of pentobarbital sodium (150 mg/kg). Animals were excluded from the final analysis in cases of surgical failure, technical issues during MRI, or unexpected death during the protocol. Final sample sizes for each experimental group are clearly indicated in the corresponding figure legends.

### Model of neonatal hypoxic-ischemic brain injury

The neonatal hypoxic-ischemic event was conducted in accordance with previously established procedures ^42^. Ischemia was induced by permanently ligating the left common carotid artery under anesthesia with 4% isoflurane for induction and 1.5% for maintenance and local lidocaine injection. Throughout the procedure, the physiological temperature of the pups was maintained constant using a rectal probe connected to a heating mat, with surgery duration not exceeding 14 minutes per pup. Following surgery, the pups were placed in a heated environment for a 30-minute recovery period. Hypoxia was induced for 2 hours in a hypoxic chamber containing 8.0 ± 0.2% O_2_ and 92% N_2_ with the chamber temperature set at 33°C to maintain a rectal temperature of the pups at 36.0 ± 0.5°C and 80% humidity. For the Sham group, only the left common carotid artery was exposed without ligation nor hypoxia- insult. After a 30-minute recovery period, the pups were kept separated from their mother in a heated environment (33 ± 1°C) for 2 hours.

### Optimal duration of hypothermia treatment

The most used hypothermia durations in the literature — 2 hours, 3 hours, and 5 hours — were tested. For all treated groups, except the HIH2Im group, pups were individually placed in a 400-ml beaker within a refrigerated water bath with circulating water, beginning 3.5 hours after ligation of the common carotid artery. The temperature was continuously monitored in additional “sentinel” pups using a rectal temperature probe. Measurements were recorded at 10-minute intervals during the first two hours of the hypothermia period and subsequently every 20 minutes. The mean rectal temperature of the rat pups was 32.3 ± 0.2°C, closely matching the target temperature of 32.0 ± 0.5°C. This target temperature was achieved within the first 20 minutes and maintained throughout the designated hypothermia durations of 2, 3, or 5 hours. For the HIH2Im group, pups were immediately individually placed in the beaker located in the refrigerated water bath for 2 hours after exiting the hypoxic chamber. Following the hypothermia procedure, the animals were rewarmed in a controlled heated environment until their body temperature reached 36°C, after which they were returned to their dams. In the normothermia condition (HIN and Sham groups), pups were placed individually in a 400-ml beaker within a water bath at 36.5°C (with rectal temperature monitored; targeted temperature was 36.0 ± 0.5°C) for 2 hours before being returned to their dams. The temperature was continuously monitored as described above. The rectal temperature for normothermia was determined based on serial temperature measurements from P7 nesting pups and was maintained at 35.8 ± 0.3°C (Fig. 2). To reduce litter-related bias, pups from each litter were randomly assigned to the different experimental groups, so that multiple conditions were represented within the same litter whenever possible.

**Figure 2:**
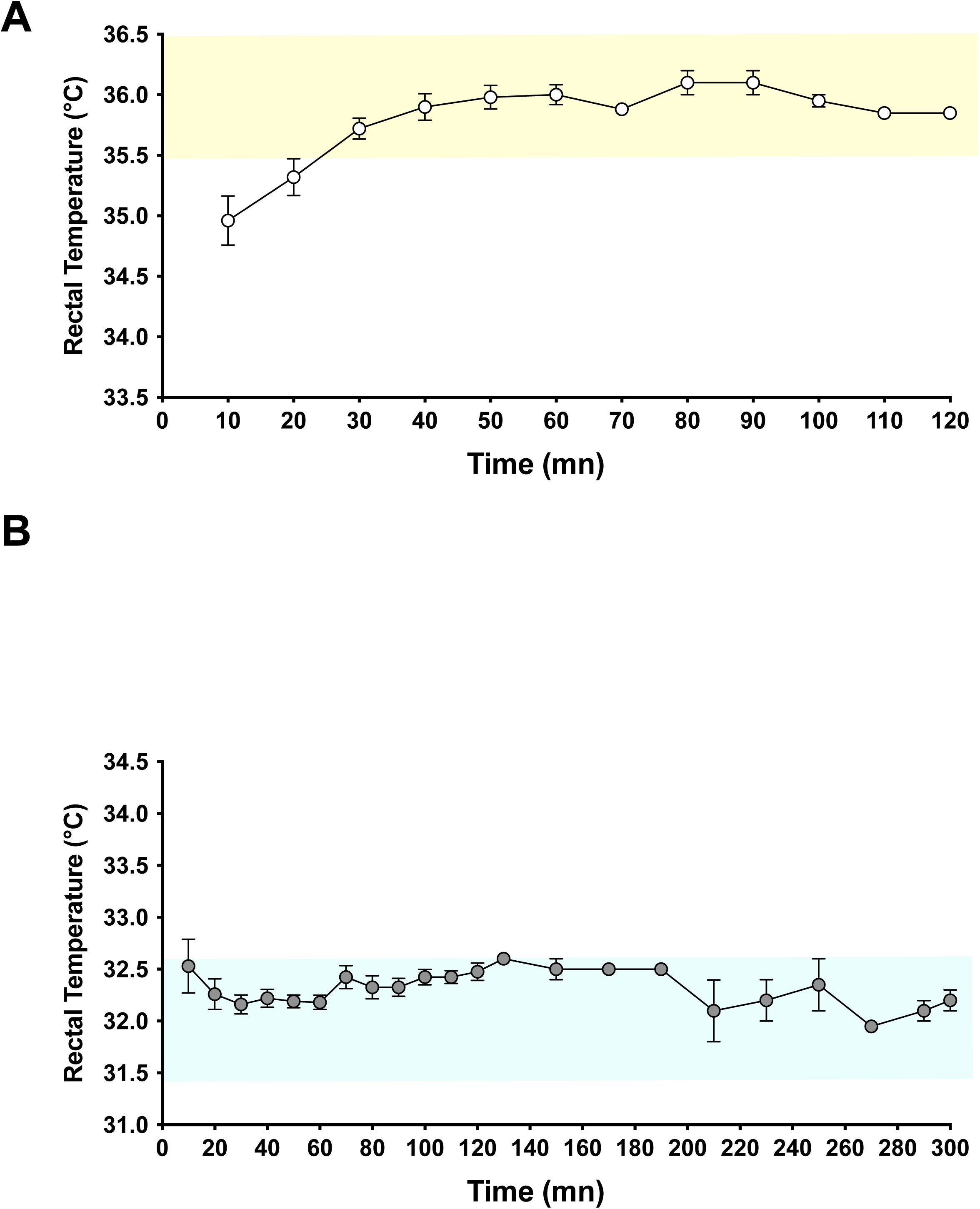
**Monitoring of rectal temperature** during hypothermia (A) or normothermia (B) procedure.

### *In vivo* MRI acquisitions and analysis

MRI scans were conducted using a horizontal 4.7T Biospec 47/50 system (Bruker, Ettlingen, Germany) equipped with a 12-cm BGA12S gradient system (660 mT/m) and acquisitions were performed at P7, P9 and P30, as previously described ^43^. Throughout all acquisitions, the pups were anesthetized with isoflurane (4% for induction and 1.5% for maintenance). Respiration was monitored using a ventral pressure sensor, and body temperature was maintained at 36.0°C using a water-heated MRI bed. Anatomical T2-weighted images of the brain were obtained using a Rapid Acquisition with Relaxation Enhancement (RARE) sequence (20 axial slices, 0.7 mm thick, echo time (TE) 50 ms, repetition time (TR) 3000 ms, total duration 4 min 48 s). Diffusion-weighted imaging (DWI) was performed with the following parameters: 20 continuous axial slices of 0.7 mm, TE 24 ms, TR 2 s, b-value 1000 s/mm², 30 directions, Δ = 8.11 ms, δ = 2.5 ms, total duration 17 min 04 s; Apparent Diffusion Coefficient (ADC) maps for each slice were computed. The first DWI scan was systematically acquired 3 hours after the common carotid artery ligation at P7. Post-processing was performed using Paravision 6.0.1 software (Bruker BioSpin, Karlsruhe, Germany). Lesion volumes were measured at P7 and P9 based on ADC maps issued from the DWI series, and at P30 based on T2-weighted slices. For each rat and each slice, and across the 20 adjacent slices per rat, regions of interest (ROI) were manually delineated to encompass the global brain area and injured area. Volumes were then calculated by considering the thickness of each slice (0.7 mm). Lesion volumes were expressed as a percentage of the total brain volume. The biological significance of lesion volume differs depending on the time point assessed. At P7, lesion volume reflects the acutely ischemic region, which includes both potentially salvageable and irreversibly damage tissue. At P9, the lesion primarily represents the infarct core, tissue that will evolve toward irreversible injury, after partial recovery or progression of the initial ischemic zone. At P30, lesion volume reflects long term structural consequences of the initial insult, including tissue necrosis, parenchymal loss and secondary atrophy (ventricular enlargement and cortical thinning). Thus, lesion volume at this late stage should be interpreted as an indicator of brain tissue loss rather than acute infarction.

### Behavioral tests

The cognitive-sensory-motor functions of pups were evaluated through a series of behavioral tests conducted from P8 to P51 ^42–45^.

#### Righting reflex

This test evaluated early motor coordination in pups (P8, P10, P12). They were placed on their backs, and the time taken to turn over onto their 4 paws was recorded in seconds (sec). Each pup underwent three trials, and the mean time to complete the reflex was calculated.

#### Modified Neurological Severity score (mNSS)

Neurological function was assessed in rats using a series of sensorimotor tests, including motor, sensory, reflex, and balance tests (P24). Each rat was individually tested, and performance deficits were assigned penalty points. The total score reflected the degree of neurological impairment: mNSS < 1: no impairment; 1–6: mild impairment; 7–12: moderate impairment and >13: severe impairment.

#### Novel object recognition

Long-term, hippocampal-dependent memory was evaluated using the novel object recognition test, conducted over a period of 4 days (P42-P45). On the first day, a 5-minute habituation period was conducted in an open field measuring 50 cm × 50 cm. On the second and third days, pups freely explored two identical objects for 5 minutes each day. On the final day, one of the familiar objects was replaced with a visually distinct new object, and pups were allowed to freely explore for 5 minutes. This procedure was recorded. Three evaluation indices were defined: the discrimination index, which measured the additional time the animal spent exploring the new object; ranging from −1 (the animal only explores the old object) to 1 (the animal only explores the new object). It was calculated as: (time spent exploring the new object - time spent exploring the familiar object) / (time spent exploring the new object + time spent exploring the familiar object); the recognition index, which evaluated the proportion of time the animal spent on the new object, calculated as: time spent exploring the new object / (time spent exploring new the object + time spent exploring the familiar object) and the exploration index, which measured the total exploration time, calculated as: time spent exploring new the object + time spent exploring the familiar object.

#### Food deprivation test

The depressive state of adult rats was evaluated using a food deprivation test conducted over four days (P48-P51). On the first two days, food rations were reduced by 20%, followed by an 80% reduction on the third day. On the fourth day, the rats were placed in an open field with black flooring and walls, with a Petri dish containing pellets soaked in 50% sucrose water placed at the center. The following parameters were measured: (1) the frequency of the animal standing on its hind legs without touching the edges; (2) the frequency of the animal sniffing and touching the food without eating; (3) the time taken to consume the pellets. The test concluded once the animal consumed the pellets. Bruxism was also recorded as a potential indicator of stress-related behavior.

### Brain removal for histology and immunohistochemistry

At P9, 19 pups were deeply anesthetized with a Ketamine-Xylazine mixture. An intracardiac perfusion with phosphate buffer saline (PBS) was performed for 10 minutes (50 ml), followed by a perfusion with paraformaldehyde (PFA) for an additional 10 minutes (50 ml), as previously described ^43^. The brains were extracted and incubated overnight in PFA at 4°C, then transferred to 30% sucrose in PBS 1X for 48 hours at 4 °C. The brains were subsequently frozen with nitrogen vapors and stored at −80 °C until sectioning with a cryostat. Histological analyses were performed on animals whose lesion volumes, assessed by MRI at P7, were within the average range of their respective experimental groups. This selection criterion was used to ensure representative sampling for immunohistological evaluation, while also allowing us to follow as many animals as possible longitudinally. Our objective was to limit the number of animals euthanized at early time points in order to preserve the cohort for later-stage behavioral and imaging assessments.

### Nissl Staining

Nissl staining was performed to estimate and compare cell death between the different conditions as previously described ^44^. In summary, 16-μm-thick brain sections at P9 were stained with Cresyl violet (0.5%, Sigma-Aldrich, France) for 10 minutes and washed in distilled water for 10 seconds. The sections were then dehydrated in 70%, 95% and 100% ethanol (2 minutes each), defatted in xylene, and mounted on coverslips. Only intact neurons with a clearly defined cell body and nuclei were quantified. Quantification was performed independently by two examiners, and the results were compared to minimize bias.

### Immunohistochemistry

To evaluate cellular degeneration, microglia activation, and polarization, 16-μm cryostat sections from the hippocampus and cortex were used. After thawing the sections at 37°C for 15 minutes and washing them in PBS six times over 30 minutes, the sections were incubated in blocking buffer (PBS containing 3% BSA and 0.3% Triton X-100, Sigma) for 1.5 hours at room temperature (RT). Subsequently, sections were incubated overnight at 4°C with primary antibodies (Table 1) in blocking buffer. After incubation, the sections were rinsed six times in PBS over 30 minutes and incubated at RT for 1 hour with secondary antibodies conjugated to Alexa Fluor, directed against goat (1:500; Alexa Fluor 488, #Ab150129) and rabbit (1:500; Alexa Fluor 568, #Ab175470). Following six additional washes in PBS, slides were mounted using VECTASHIELD mounting medium containing 4′,6-diamidino-2-phenylindole (DAPI) (Vector Laboratories, Eurobio, France) for nuclear staining. Images were captured using 20x and 40x objective with an Eclipse 90i microscope (Nikon).

**Table 1.** Primary antibodies used for staining.

| Host/ Primary antibody | Producer | Antibody type | Dilution |
| --- | --- | --- | --- |
| Goat/ Anti-IBA1 | Abcam | Monoclonal | 1:1000 |
| Rabbit/Anti-CD86 | Proteintech | Polyclonal | 1:4000 |
| Rabbit/ Anti-Mannose receptor (CD206) | Abcam | Polyclonal | 1:10000 |
| Rabbit/ Anti-Caspase 3 | Abcam | Polyclonal | 1:500 |

### Quantification of cell loss, microglia activation and polarization

Image acquisition of the brain sections was performed using NIS Elements 4.30 software, and image processing was conducted with ImageJ software. Initially, three fluorescence images were obtained: the first in the DAPI fluorescence channel, the second in the FITC fluorescence channel, and the third in the Cyanine 3 fluorescence channel. Cellular loss was evaluated in Caspase3-stained tissue sections, while microglia activation and polarization were assessed in tissue sections stained for Iba1 and CD86/CD206. Quantification of fluorescence intensity for each marker was performed by averaging measurements obtained from two brain sections per animal, taken from the ipsilateral hemisphere. Analyses were specifically carried out in the CA1 subfield of the hippocampus and in the cortical region. The average values for each condition were then calculated using the following equation to determine the corrected total cell fluorescence (CTCF). CTCF = Integrated density – (Area of selected cell x Mean fluorescence of background readings). Fluorescence images from all samples were independently quantified by a blind examiner.

### Microglial Morphology Analysis

Microglial morphology was assessed using Iba1 immunofluorescence images. Images were first opened and converted to 8-bit format in ImageJ software. A threshold was applied to distinguish microglial structures from the background, and images were subsequently binarized and skeletonized using the “Binary Skeleton” tool. To remove small artifacts, the “Analyze Particles” function was applied with a size range of 50 pixels to infinity. Skeletonized images were then analyzed using the Skeleton Analysis plugin to quantify morphological parameters, including the number of branches, junctions, and triple points. All measurements were performed under consistent image processing settings to ensure comparability between conditions.

### Statistical analysis

Except for bruxism and mNSS, all data are presented as mean ± SD. For bruxism, data are presented as mean with range. Data from mNSS were reported as medians with interquartile ranges (IQR). Statistical analyses were conducted using GraphPad Prism V10. Prior to statistical testing, data normality was assessed using the Shapiro–Wilk test. For datasets exhibiting a normal distribution, one-way analysis of variance (ANOVA) was used for multiple-group comparisons, followed by Fischer’s LSD post hoc test. For datasets that did not meet normality assumptions, the Kruskal–Wallis test was applied, followed by Dunn’s multiple-comparisons test. Comparisons between the HIH2 and HIH2Im groups, as well as between males and females within each group, were performed using the unpaired Student’s t-test. When normality assumptions were not met, the Mann-Whitney test was used instead. A p-value of < 0.05 was considered statistically significant.

## RESULTS

### Mortality rate

Of the 77 animals (both male and female) included in this study, 13 were not subjected to hypoxia-ischemia (HI) and demonstrated a 100% survival rate. Among the 64 animals exposed to the hypoxic-ischemic procedure, 10 (6 males and 4 females) did not survive, resulting in a mortality rate of 15.6%.

### Optimal hypothermia duration for short-term neuroprotection

The short-term neuroprotection provided by varying durations of therapeutic hypothermia was assessed using diffusion MRI to monitor cerebral lesion volume and by evaluating the righting reflex for functional assessment in newborns (Fig. 3). Diffusion MRI was performed at P7 and P9 and brain lesions volumes were measured on DWI, seen as diffusion restriction area (Fig. 3A). At P7, the day of the hypoxic-ischemic event, no pups have yet been treated with therapeutic hypothermia (as therapeutic hypothermia began after the MRI acquisitions); therefore, no significant differences were found at this time point (Fig. 3B). At P9, pups in the HIH2 group exhibited significantly smaller brain lesion volumes compared to those in the HIN group (29 ± 6% vs. 39 ± 7%, respectively). No significant differences in the brain lesion volumes were observed between the HIN group and the other therapeutic hypothermia durations (HIN vs. HIH3 and HIH5; Fig. 3B). The reduction in lesion volume 48 hours after the hypoxic-ischemic insult was calculated using the following formula: ((Lesion volume at P7−Lesion volume at P9)/ (Lesion volume at P7) × 100). Interestingly, pups in the HIH2 group exhibited a greater lesion volume reduction compared to those in the other groups, with percentage reductions of 33 ± 10% for the HIH2 group, 23 ± 9% for HIH3, 21 ± 12% for HIH5, and 17 ± 9% for the HIN group. (Fig. 3C). The righting reflex was assessed from P8 to P12 (Fig. 3D). At P8, the Sham group showed the best performance and only the pups from the HIH2 group had similar scores. At P10, among hypothermia-treated groups, only HIH2 pups performed better than those in the HIN group. At P12, HIH2 group showed better performance than HIN as well as other hypothermia-treated groups, reaching scores comparable to the Sham group. Additionally, no sex-specific differences were found between any groups regarding brain lesions at P9 or in the early reflex test (Supplementary Fig. S1 and S3).Two hours of hypothermia showed the best protection on these parameters (brain lesions volumes at short term and righting reflex) compared to 3 and 5 hours.

**Figure 3:**
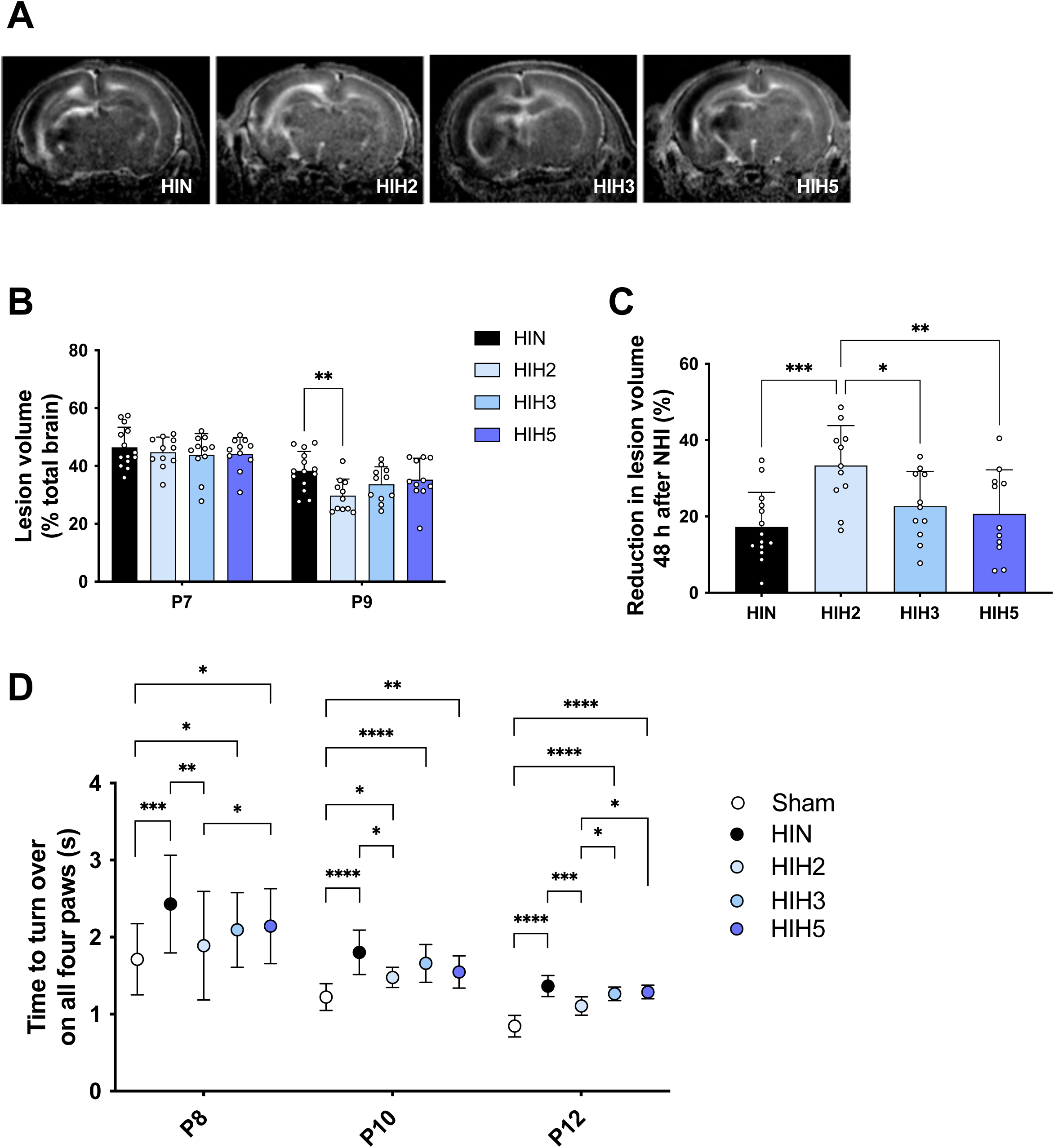
Assessment of varying durations of therapeutic hypothermia for short-term neuroprotection. (A) ADC maps from DWI of P9-pup brains acquired 3 h after the surgery for the HIN (n = 14), HIH2 (n = 11), HIH3 (n = 11), and HIH5 (n = 11) groups. (B) Quantifications of brain lesion volumes (dark signal), expressed in % relative to total brain volume, for the different groups. (C) Reduction in lesion volume (between P7 and P9) after NHI injury, express in %, for the different groups. (D) Righting reflex test at P8, P10, P12, for the different groups. * p < 0.05; ** p < 0.01; *** p < 0.001; **** p < 0.0001.

### Optimal hypothermia duration for long-term neuroprotection

T2-weighted MRI conducted at P30 enabled the assessment of long-term lesion volumes (Fig. 4A). At P30, only the HIH2 group exhibited significantly smaller lesion volumes compared with the HIN group (brain lesion volumes: 6 ± 11% vs. 17 ± 14% for HIH2 and HIN, respectively; Fig. 4B). Moreover, among hypothermia-treated groups, the HIH2 group showed significantly reduced lesion volumes compared with the HIH5 group (6 ± 11% vs. 19 ± 16% for HIH2 and HIH5, respectively). For each experimental group, no sex differences in brain lesion volumes at P30 were noted (Supplementary Fig. S2). Concerning behavioral tests (Fig. 4C), only pups in the HIH2 group showed mNSS performance comparable to the Sham group. Both HIH2 and HIH3 groups exhibited significantly improved scores compared to HIN, whereas HIH5 did not differ from HIN. No sex differences were observed in mNSS performance (Supplementary Fig. S4). No statistical differences were observed on brain lesion volumes and mNSS evaluation between HIH2 and HIH3.

**Figure 4:**
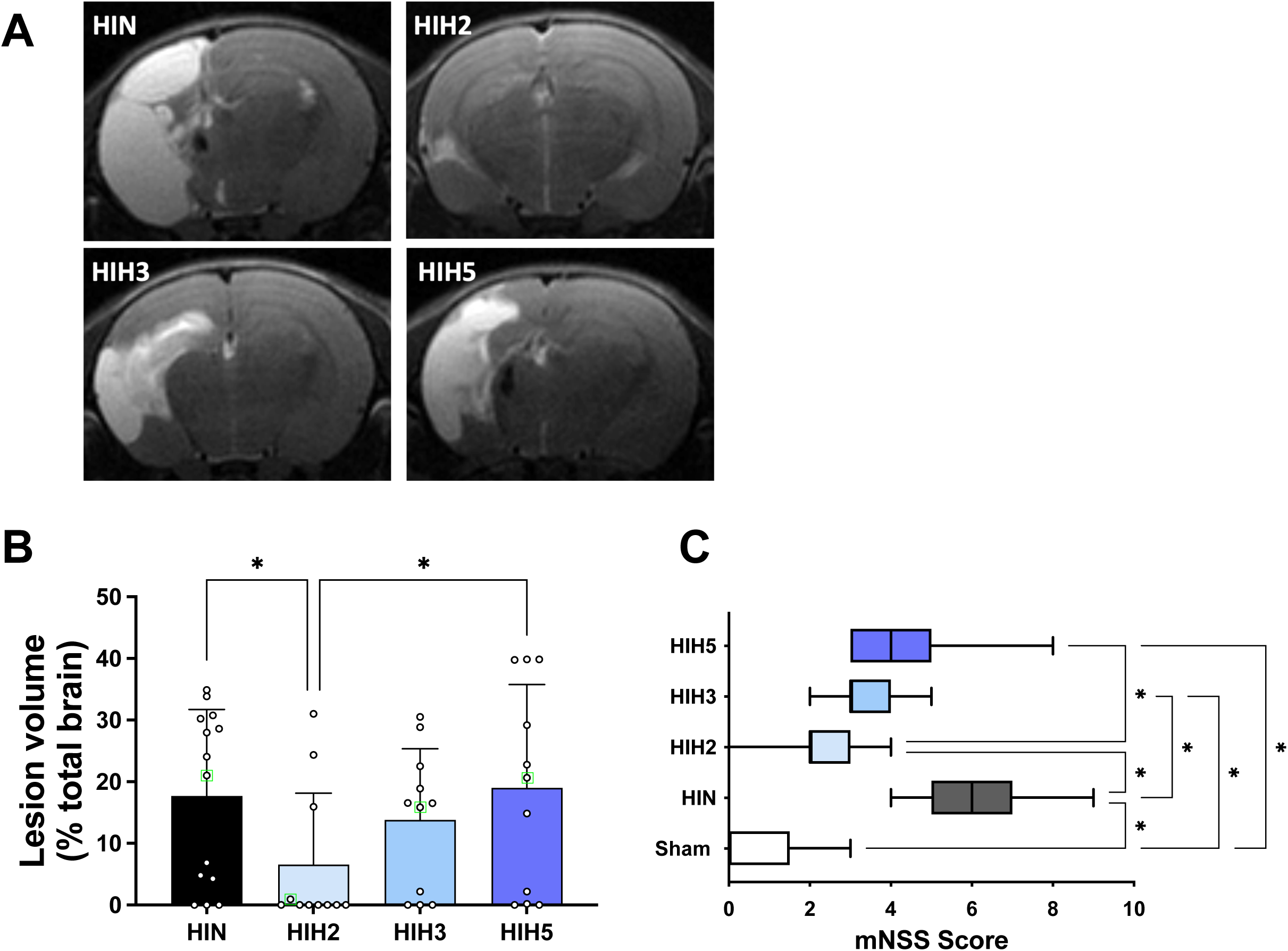
Assessment of varying durations of therapeutic hypothermia for long-term neuroprotection. (A) T2-RARE MRI of P30 pup brains acquired 23 days after the HI event for the HIN (n = 14), HIH2 (n = 11), HIH3 (n = 11), and HIH5 (n = 11) groups. (B) Quantification of brain lesion volumes, expressed as a percentage of total brain volume, for the different groups. (C) Modified Neurological Severity Scores (mNSS) at P24, for the different groups. * p < 0.05.

### Optimal hypothermia duration for cognitive function preservation

Given that NHI is associated with cognitive impairment, the neuroprotective effects of therapeutic hypothermia on long-term memory were initially assessed using the novel object recognition test at P45 (Fig. 5). The discrimination index was the lowest in the HIN group compared to the Sham group (0.05 ± 0.20 vs.0.90 ± 0.05, respectively). In HIH2 and HIH3 and HIH5 groups, the discrimination index was higher, compared to HIN (0.70 ± 0.12; 0.60 ± 0.26, 0.35 ± 0.24 and 0.05 ± 0.02, for HIH2, HIH3, HIH5 and HIN groups, respectively, Fig. 5A). In addition, the discrimination index of the HIH2 and HIH3 groups were significantly higher than that of the HIH5 group. No sex differences were found between experimental groups regarding the discrimination index (Supplementary Fig. S5). The Sham group exhibited also the highest recognition index (0.95 ± 0.03), indicating optimal cognitive performance (Fig. 5B). In contrast, the lowest values were observed in the HIN (0.55 ± 0.04) and HIH5 (0.62 ± 0.17) groups. Intermediate improvements were observed in the HIH2 and HIH3 groups (recognition index: 0.84 ± 0.06 and 0.77 ± 0.15, respectively). Finally, the highest exploration indices were recorded in both the Sham and HIH2 groups, which did not differ statistically, and were significantly higher than those observed in the HIN, HIH3, and HIH5 groups (Fig. 5C).

**Figure 5:**
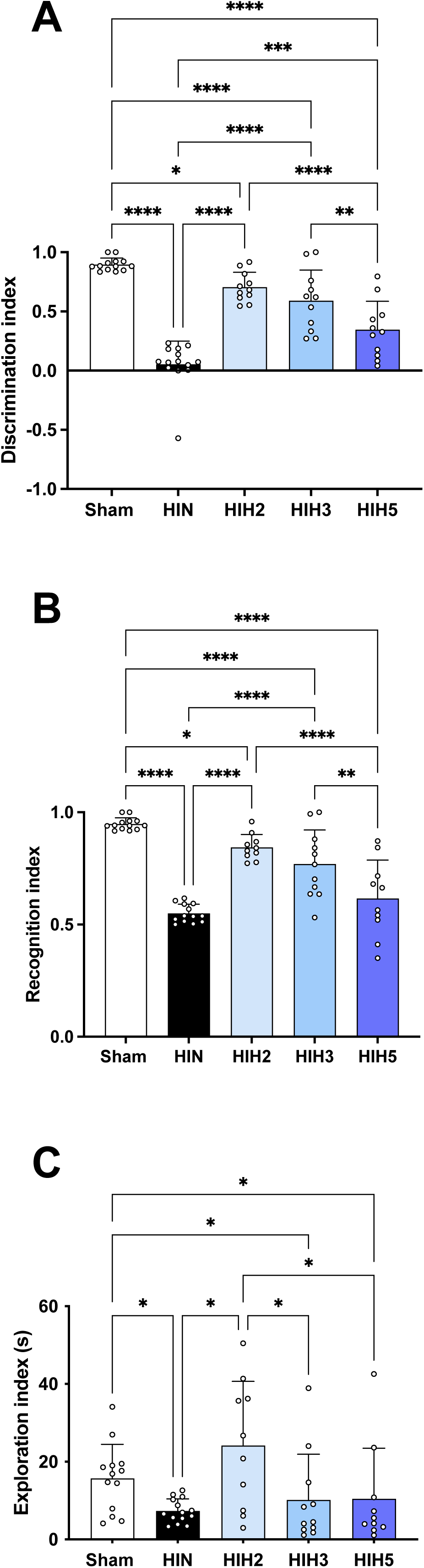
Assessment of varying durations of therapeutic hypothermia on long-term memory impairment (P45), for Sham (n = 13), HIN (n = 14), HIH2 (n = 11), HIH3 (n = 11), and HIH5 (n = 11) groups. (A) Discrimination index. (B) Recognition index. (C) Exploration index. * p < 0.05; ** p < 0.01; *** p < 0.001; **** p < 0.0001.

Since emotional regulation may also be affected during adolescence following a neonatal hypoxic-ischemic event, the depressive and anxious states of the rats were assessed at postnatal day 51 using the food restriction test (Fig. 6). Therapeutic hypothermia for two hours (HIH2) allowed pups to show similar latency to eat as the Sham group after food restriction, unlike the HIN, HIH3, and HIH5 groups (Fig. 6A). Latency times were 147 ± 43 s (Sham), 200 ± 117 s (HIH2), 408 ± 192 s (HIN), 363 ± 198 s (HIH3), and 438 ± 226 s (HIH5). Additionally, only the Sham and HIH2 groups showed significantly more frequent engagement with food, either by smelling or touching it, compared to the HIN and HIH5 groups (Fig. 6B). Furthermore, during the test, 46% of the HIN rats exhibited bruxism, a sign of stress or anxiety, compared to 0%, 9%, 18%, and 27% in the Sham, HIH2, HIH3, and HIH5 groups, respectively (Fig. 6C). Again, only pups in the Sham and HIH2 groups differed significantly from those in the HIN group. However, regarding these depressive and anxiety-like behaviors, no differences were observed between the HIH2 and HIH3 groups. No sex-specific differences were observed across the groups (Supplementary Fig. S6).

**Figure 6:**
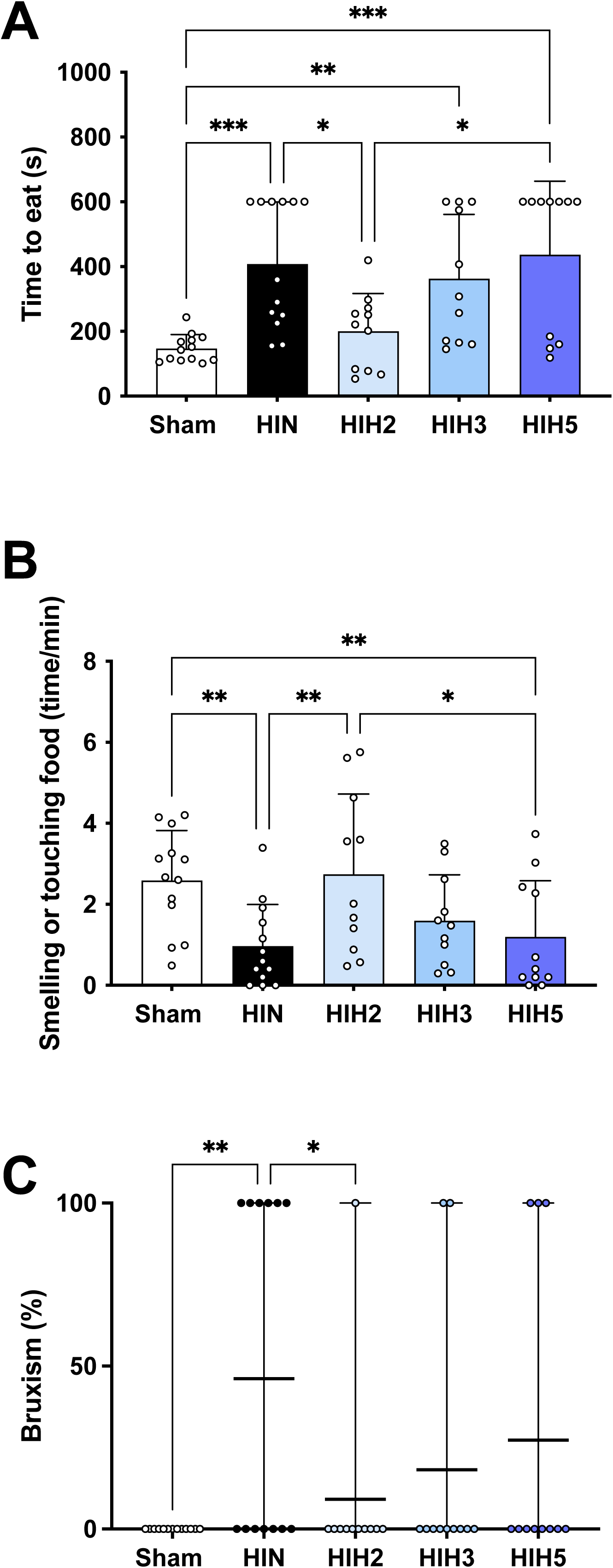
Assessment of varying durations of therapeutic hypothermia on anxiety and depressive status induced by neonatal hypoxia-ischemia, for Sham (n = 13), HIN (n = 14), HIH2 (n = 11), HIH3 (n = 11), and HIH5 (n = 11) groups. (A) Time to eat the pellet during the food restriction test. (B) Time spent to smell or touch food during the test. (C) Proportion of bruxism (%). * p < 0.05; ** p < 0.01; *** p < 0.001.

### Effect of the hypothermia duration on neuronal death

The protective effect of hypothermia on neuronal death was assessed using Nissl staining. In the cerebral cortex and hippocampus, cell nuclei in the Sham group displayed typical and homogenous morphologies. In the contralateral hemisphere, hypoxia-ischemia did not lead to significant cortical and hippocampal changes in Nissl staining (Supplementary Fig. S7). However, in the ipsilateral hemisphere, significant neuro-morphological alterations were observed in the HIN group, particularly in the cerebral cortex and hippocampus (CA1 area). The cellular architecture was disrupted, and a significant proportion of neurons exhibited cell death (Fig. 7A). The number of Nissl-stained nuclei was significantly reduced in the HIN group in both the injured cortex and the damaged CA1 region of the hippocampus (26 ± 6% and 39 ± 6%, respectively), relative to the Sham group, to which all values were normalized (Fig. 7B–C). All hypothermia-treated groups showed better preservation of neuronal architecture and Nissl substance in the ipsilateral cortex compared with the HIN group, with the HIH5 group also exhibiting enhanced cortical cell viability, although not reaching Sham levels; in the CA1 region, this protective effect was observed only in the 2- and 3-hour hypothermia groups (Fig. 7B–C).

**Figure 7:**
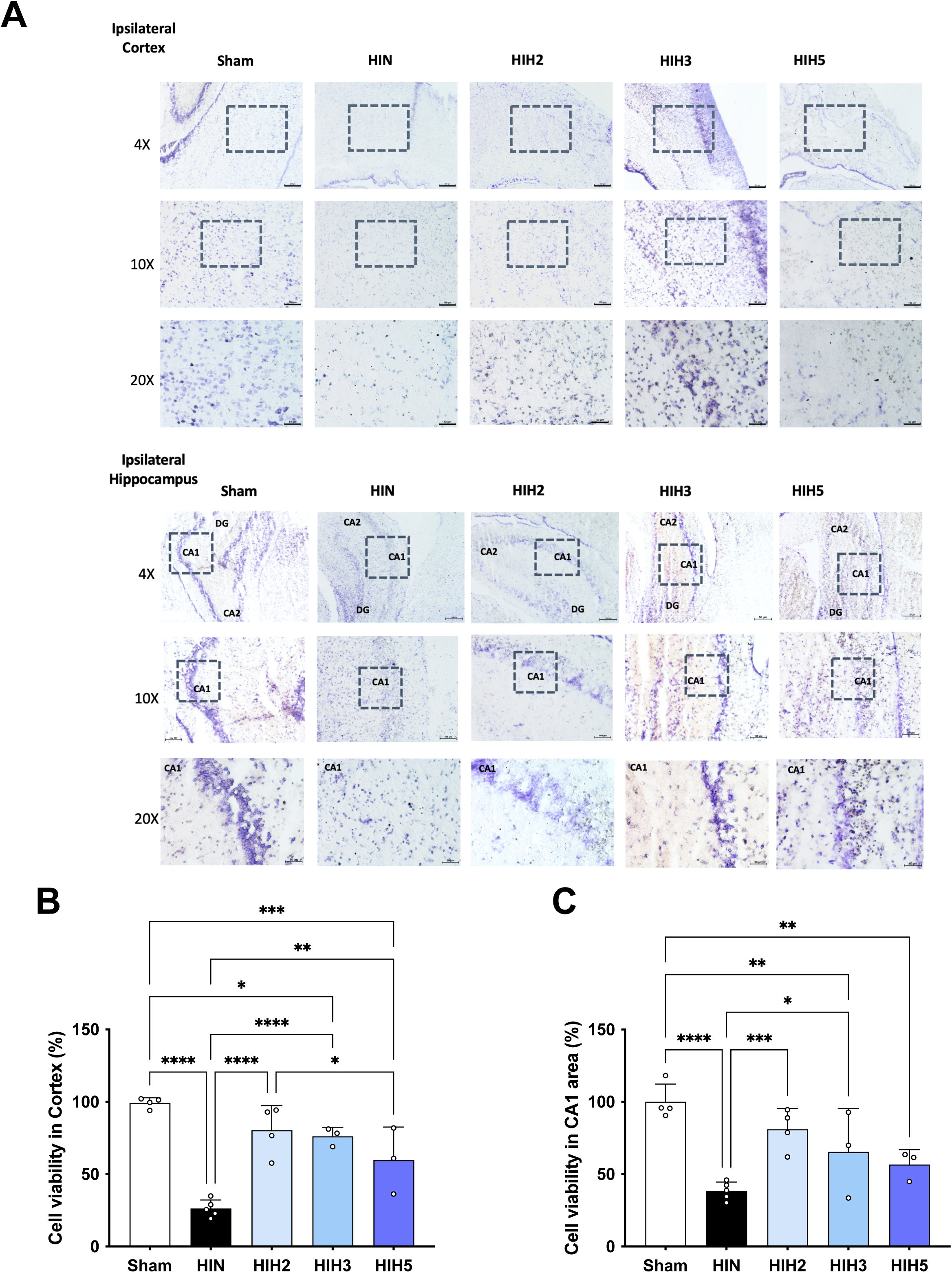
Histology 48 h after neonatal hypoxia-ischemia. (A) Representative image of Nissl staining in the cortex and in the hippocampus CA1 regions. Scale bars represent 250 μm in the images of 4X; 100μm in the images of 10X; 50μm in the images of 20X and 20μm in the images of 40X. (B) Percentage of cell loss, based on the density of Nissl-stained nuclei, for Sham (n = 4; 2 fields per brain), HIN (n = 5; 2 fields per brain), HIH2 (n = 4; 2 fields per brain), HIH3 (n = 3; 2 fields per brain) and HIH5 (n = 3; 2 fields per brain) groups in ipsilateral cortex). (C) Percentage of cell loss, based on the density of Nissl-stained nuclei, for Sham (n = 4; 2 fields per brain), HIN (n = 5; 2 fields per brain), HIH2 (n = 4; 2 fields per brain), HIH3 (n = 3; 2 fields per brain) and HIH5 (n = 3; 2 fields per brain) groups in hippocampus CA1 area. * p < 0.05; ** p < 0.01; *** p < 0.001; **** p < 0.0001.

### Effect of hypothermia on microglial activation and apoptosis

Because 2-hour hypothermia appeared slightly more effective than 3-hour treatment, immunohistochemical analyses were performed to investigate at the cellular and molecular levels whether 2 h provides superior effects compared with 3 h in the HIN, HIH2, HIH3, HIH5, and Sham groups. First, the impact of hypothermia treatment was assessed on overall microglia activation by performing immunohistochemistry for Iba-1 in the ipsilateral cortex and hippocampus (CA1 area) (Fig. 8). The HIN group exhibited a significant increase in Iba-1 fluorescence intensity in both regions compared with the Sham group (Fig. 8A-B). Among the hypothermia-treated groups, the HIH2 and HIH3 groups showed reduced Iba-1 fluorescence relative to the HIN group in both regions. Moreover, the HIH2 group displayed significantly lower Iba-1 fluorescence than the HIH5 group in both regions. No significant differences in Iba-1 fluorescence were observed between the HIH2 and HIH3 groups in either region. To further characterize microglial activation states, morphological complexity was quantified using skeleton analysis (Fig. 8C). In the cortex and the CA1 region of the hippocampus, neonatal hypoxia-ischemia (NHI) induced a pronounced simplification of microglial arborization, evidenced by a significant reduction in the number of branches compared to Sham controls (Fig. 8C). Notably, among the hypothermia-treated groups, only the 2-hour hypothermia condition restored the number of branches to levels comparable with Sham animals. Similarly, the number of junctions, reflecting the complexity of branching architecture, was decreased following NHI and returned to Sham levels in the HIH2 group (Supplementary Fig. S8). Consistent with these observations, NHI also reduced the number of branching points (Supplementary Fig. S8), whereas 2 hours of hypothermia preserved the structural complexity of microglia.

**Figure 8:**
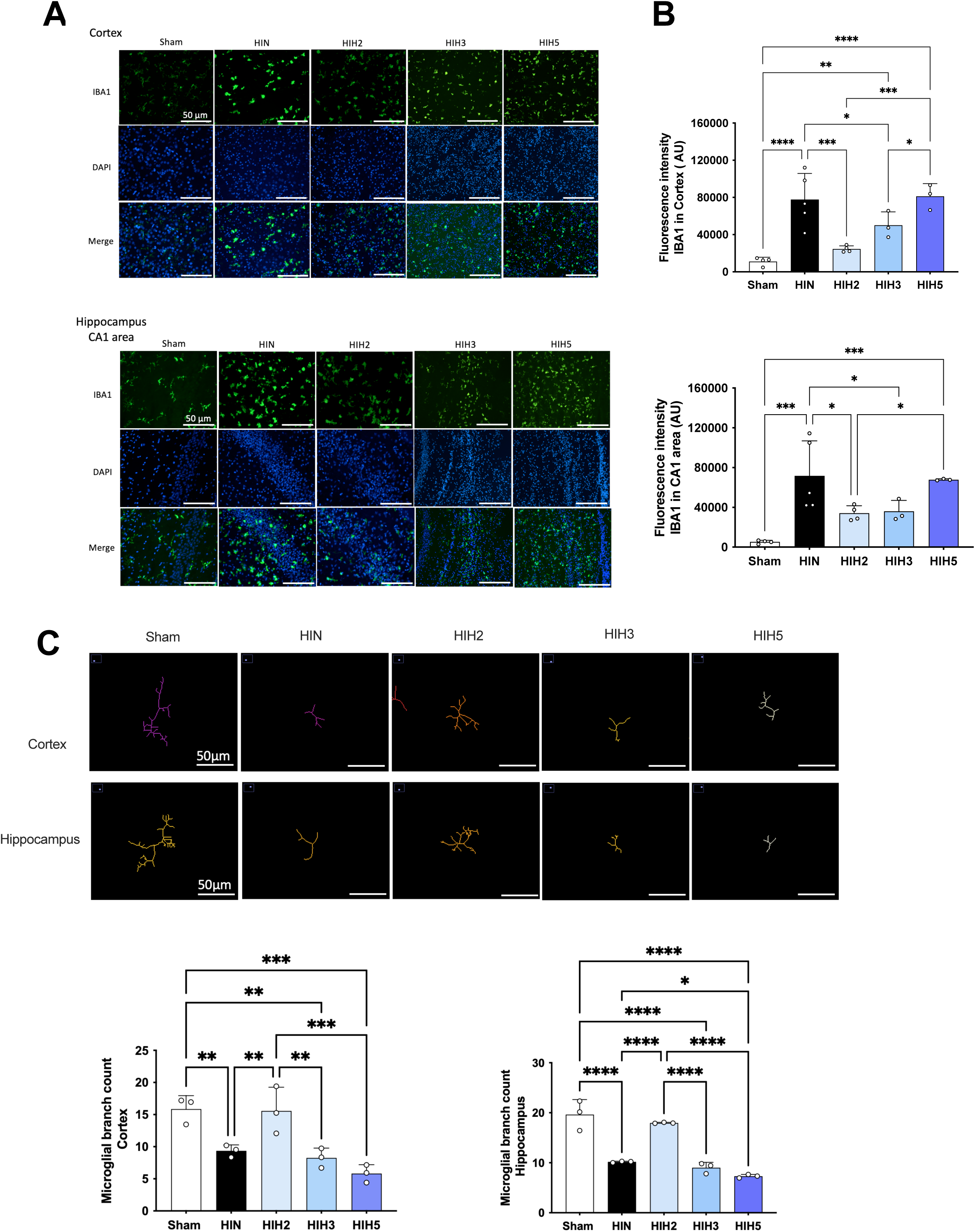
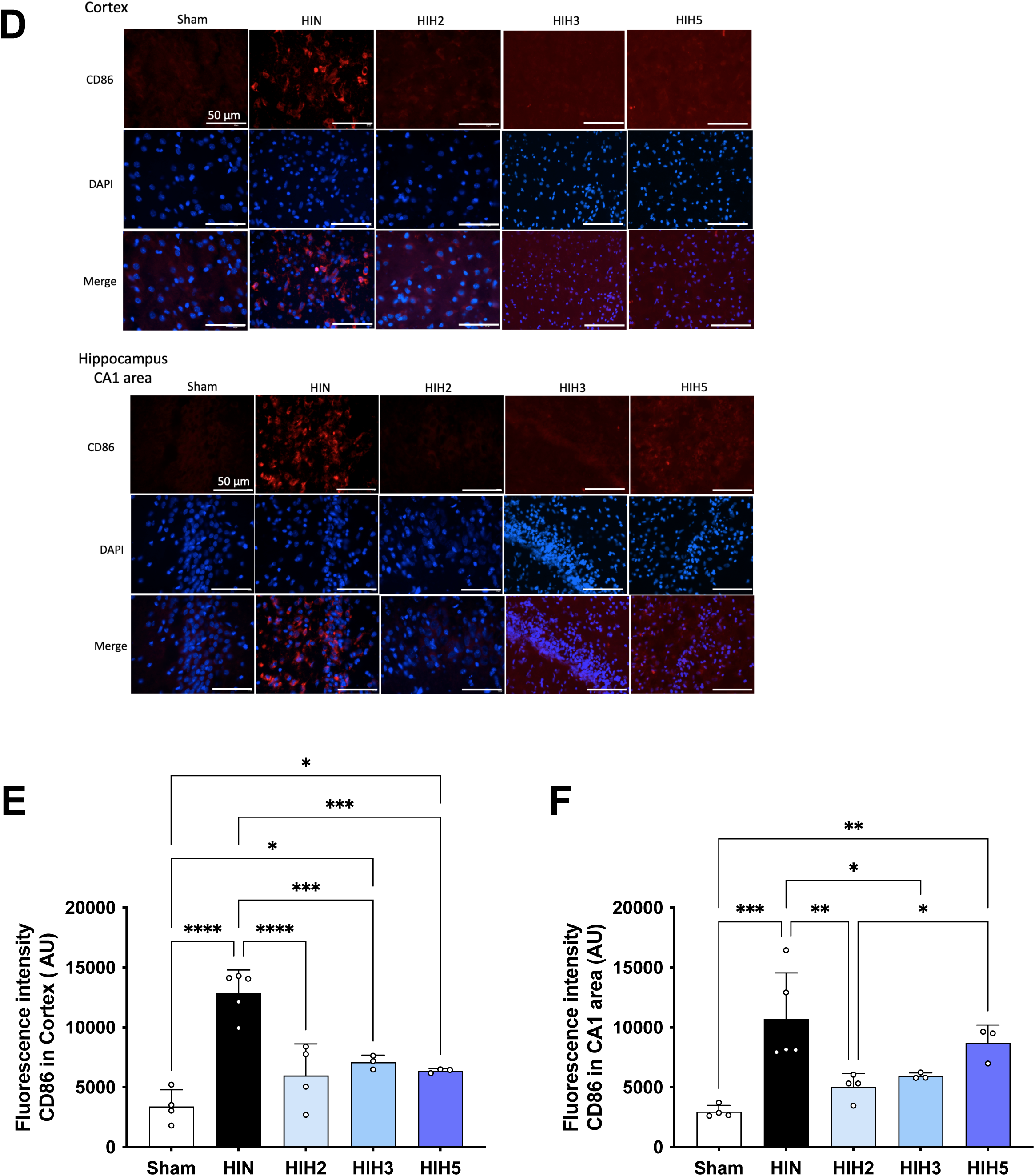
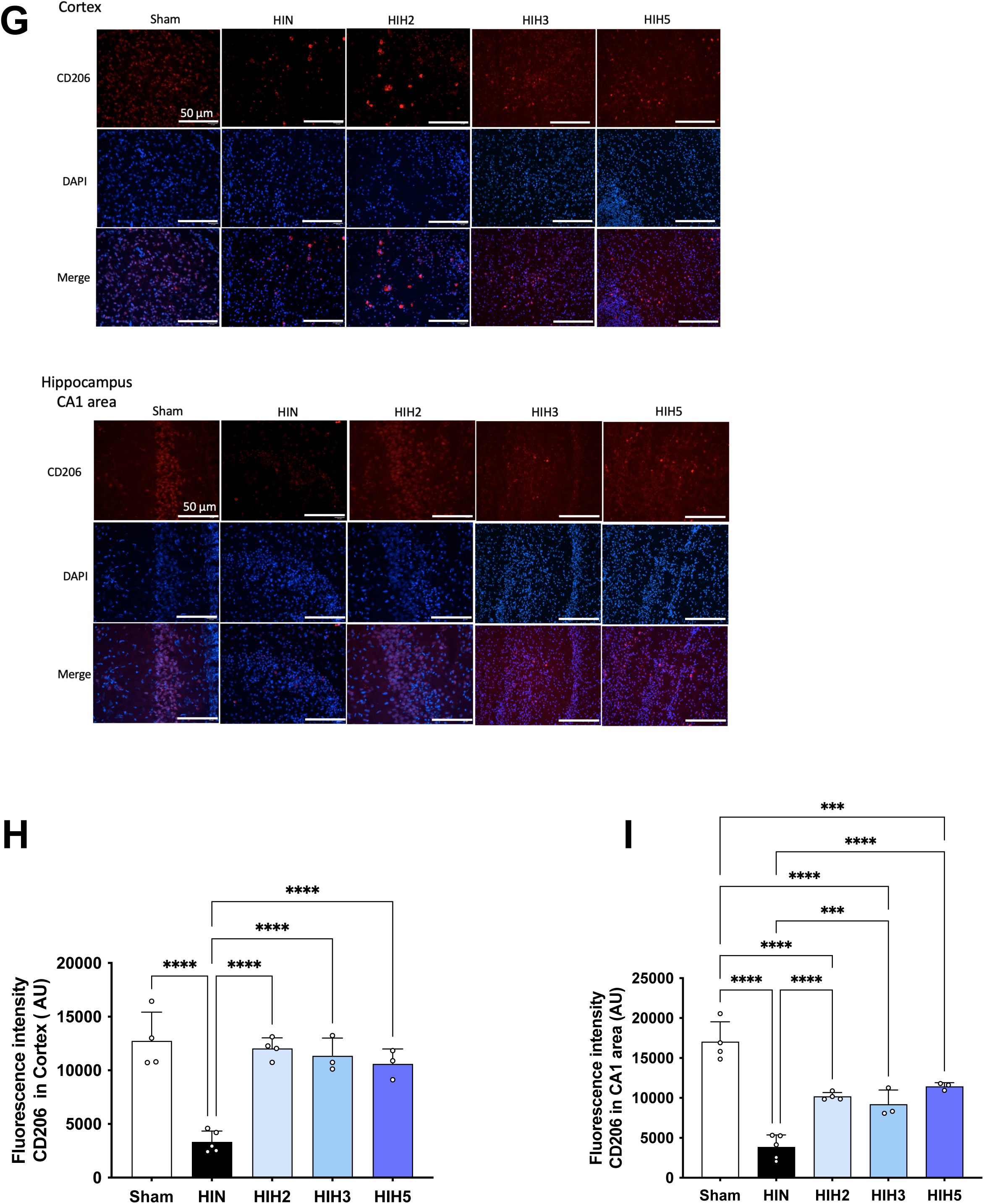
Impact of a 2h-hypothermia treatment after NHI on microglia activation. (A) Representative immunofluorescence images of Iba1 expression in the cortex and hippocampal CA1 region across experimental groups at 48 h post-NHI (20× magnification; scale bar, 50 μm). (B) Quantification of Iba1 fluorescence intensity in the cortex (top) and hippocampal CA1 region (bottom) for Sham (n = 4), HIN (n = 5), HIH2 (n = 4), HIH3 (n = 3), and HIH5 (n = 3) groups. For each brain, fluorescence intensity was averaged from two fields. (C) Top: Representative images of reconstructed microglial skeletons in the cortex and hippocampus across experimental groups. Bottom: Quantification of microglial branch number in the cortex and hippocampus. (D) Representative immunofluorescence images of CD86 expression in the cortex and hippocampal CA1 region at 48 h post-NHI (20× magnification; scale bar, 50 μm). (E,F) Quantification of CD86 fluorescence intensity in the cortex (E) and hippocampal CA1 region (F). (G) Representative immunofluorescence images of CD206 expression in the cortex and hippocampal CA1 region at 48 h post-NHI (20× magnification; scale bar, 50 μm). (H,I) Quantification of CD206 fluorescence intensity in the cortex (H) and hippocampal CA1 region (I). Data are shown for Sham (n = 4), HIN (n = 5), HIH2 (n = 4), HIH3 (n = 3), and HIH5 (n = 3) groups, with values averaged from two fields per brain. Statistical significance is indicated as *p < 0.05; **p < 0.01; ***p < 0.001; ****p < 0.0001.

To further characterize myeloid cells responses following NHI event, the expressions of CD86 and CD206, markers associated pro-inflammatory (M1-like) and anti-inflammatory (M2-like) microglial phenotypes, respectively, were evaluated in the ipsilateral cortex and hippocampus (CA1 area). CD86 fluorescence intensity was increased after HI in the HIN group, significantly higher from the intensities measured in the Sham and HIH2 groups in both regions (Fig. 8D-F). In contrast, the HIH2 and HIH3 groups exhibited reduced CD86 fluorescence in both regions. Five hours of hypothermia led to a reduction on CD86 fluorescence intensity only in the cortex, not in the hippocampus. Simultaneously, CD206 fluorescence intensity was markedly higher in all hypothermia-treated groups in both regions, and without any statistical differences with the Sham group in the cortex, while it was decreased in the HIN group in both regions (Fig. 8G-I).

Caspase-3 activation was also assessed 48 hours after NHI injury to examine the effects of NHI and hypothermia on apoptosis. Immunofluorescence staining in the hippocampus (CA1 area) and cortex showed increased fluorescent intensity in the HIN group compared to the Sham group (Fig. 9A-C). Caspase-3 signals in the cortex were reduced in all hypothermia treated-groups compared with the HIN group (Fig. 9B). Concerning the hippocampus, only the HIH2 group exhibited lower caspase-3 fluorescence comparted to the HIN group (Fig. 9C).

**Figure 9:**
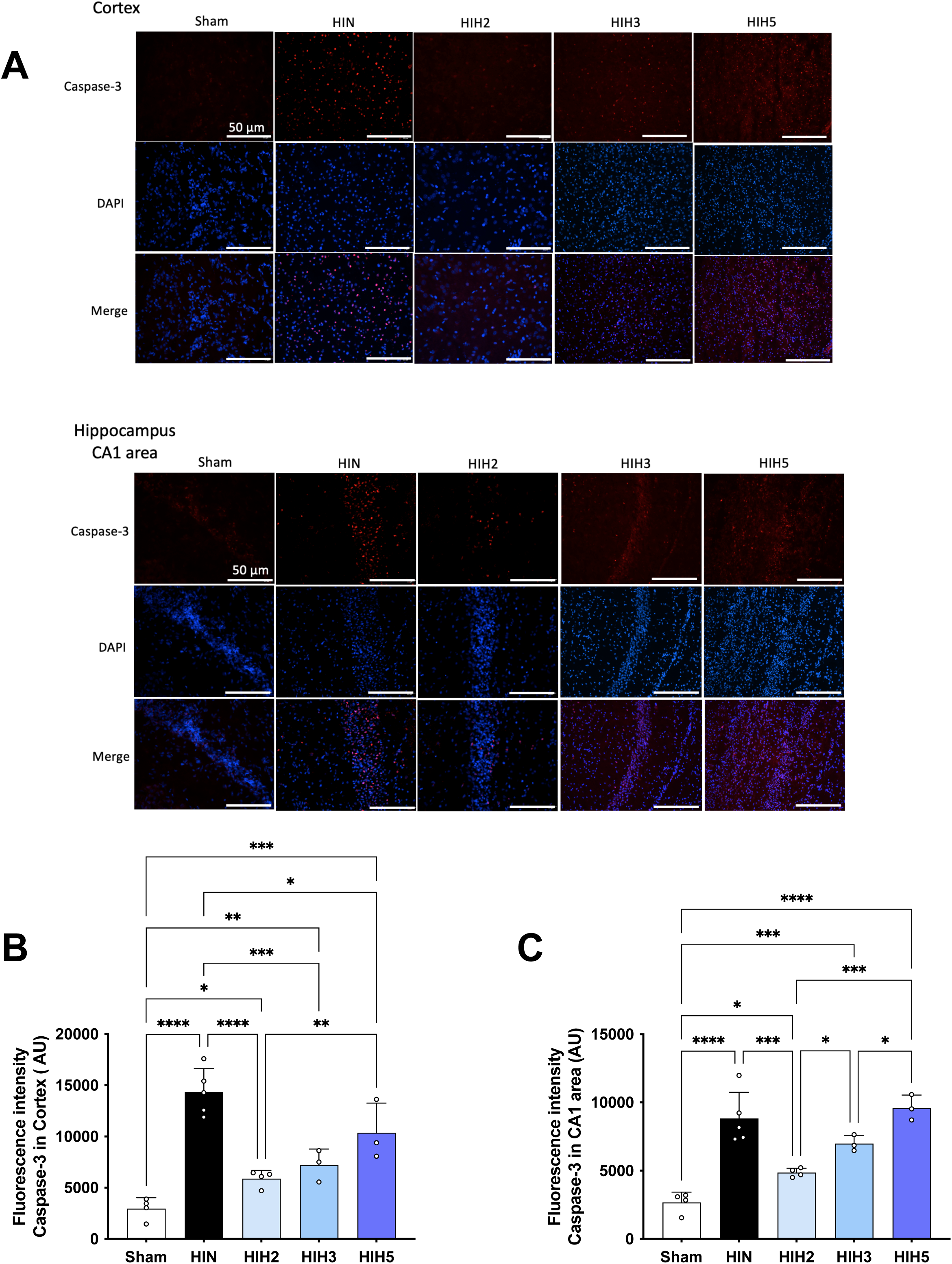
Impact of a 2h-hypothermia treatment after NHI on apoptosis. (A) Representative immunofluorescence images of Caspase-3 expression in the cortex and hippocampal CA1 region across experimental groups (20× magnification; scale bar, 50 μm). (B,C) Quantification of Caspase-3 fluorescence intensity in the cortex (B) and hippocampal CA1 region (C). Data are shown for Sham (n = 4), HIN (n = 5), HIH2 (n = 4), HIH3 (n = 3), and HIH5 (n = 3) groups, with values averaged from two fields per brain. * p < 0.05; ** p < 0.01; *** p < 0.001; **** p < 0.0001.

### Effect of timing on the efficacy of therapeutic hypothermia

As a final point, we wondered whether the timing of hypothermia initiation could influence the extent of neuroprotection. To explore this, data from the HIH2 group were compared with those from the HIH2Im group, in which therapeutic hypothermia was initiated immediately upon removal from the hypoxic chamber, therefore 1 ½ hour before the HIH2 group (Fig. 10). At P9, brain lesion volumes measured by MRI were smaller in the HIH2Im group compared to the HIH2 group (29 ± 6% and 21 ± 6% for the HIH2 and HIH2Im groups, respectively) (Fig. 10A). However, at P30, no significant differences in lesion volume were observed between the two groups. No differences were observed between the two groups in the righting reflex test or the neurological score (mNSS) (Fig. 10B-C). Nevertheless, in the novel object recognition test, the HIH2Im group exhibited significantly higher discrimination and recognition indices compared to the HIH2 group (Fig. 10D-E), while the exploration index remained similar between groups (Fig. 10F). Additionally, no significant differences were found between groups regarding the impact of hypothermia initiation timing on depressive-like behavior in rats (Fig. 10G-H).

**Figure 10:**
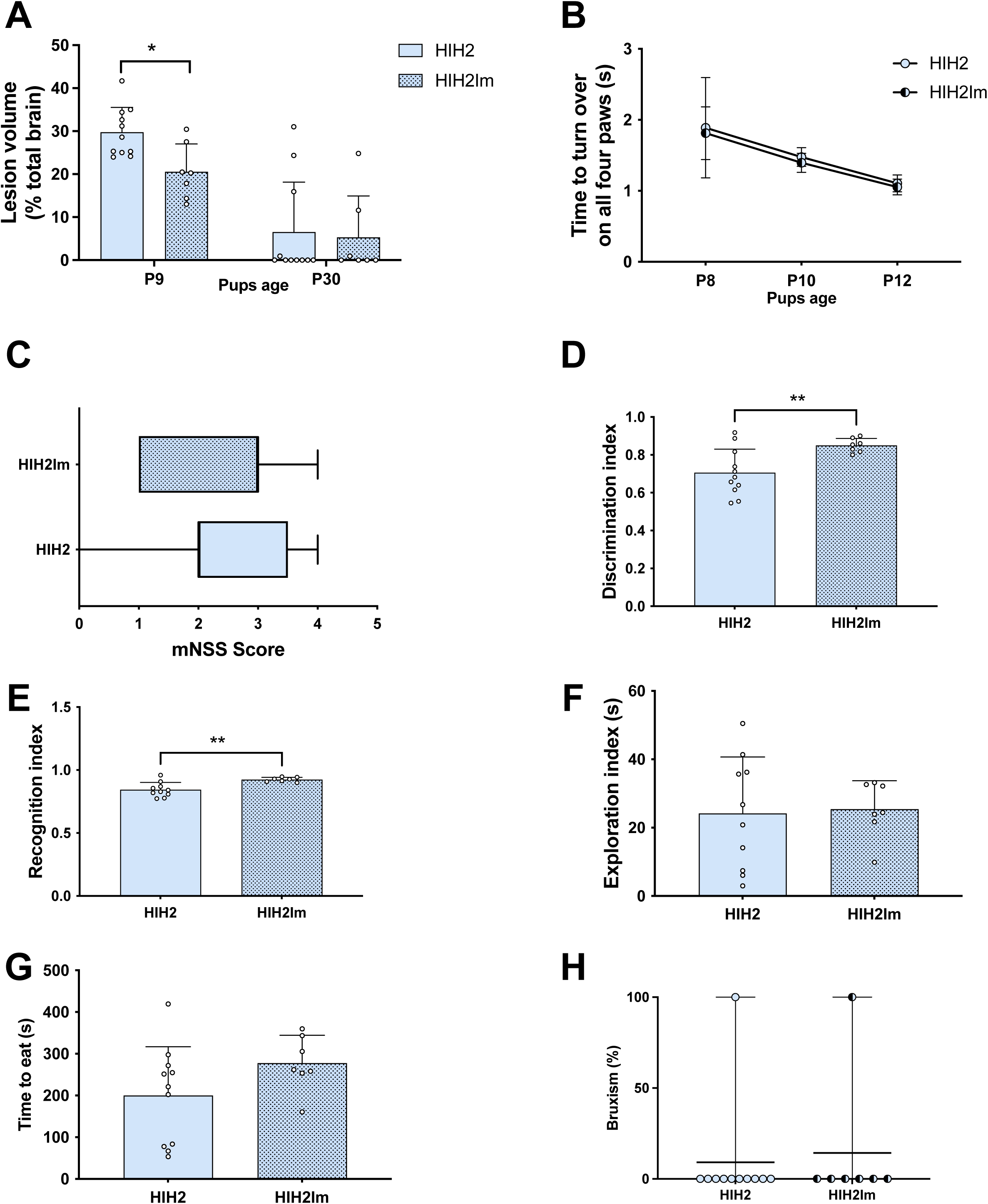
Effects of delayed versus immediate initiation of therapeutic hypothermia on brain lesion volume and behavioral outcomes. (A) Quantifications of brain lesion volumes, at P9 and P30, express in % relative to total brain volume, for the different groups. (B) Righting reflex test at P8, P10, P12, for the different groups. (C) mNSS at P24, for the different groups. (D) Novel object recognition, discrimination index. (E) Novel object recognition, recognition index. (F) Novel object recognition, exploration index. (G) Time to eat the pellet during the food restrictive test. (H) Proportion of bruxism. * p < 0.05 and ** p < 0.01.

A summary of the data is presented Table 2.

**Table 2.**
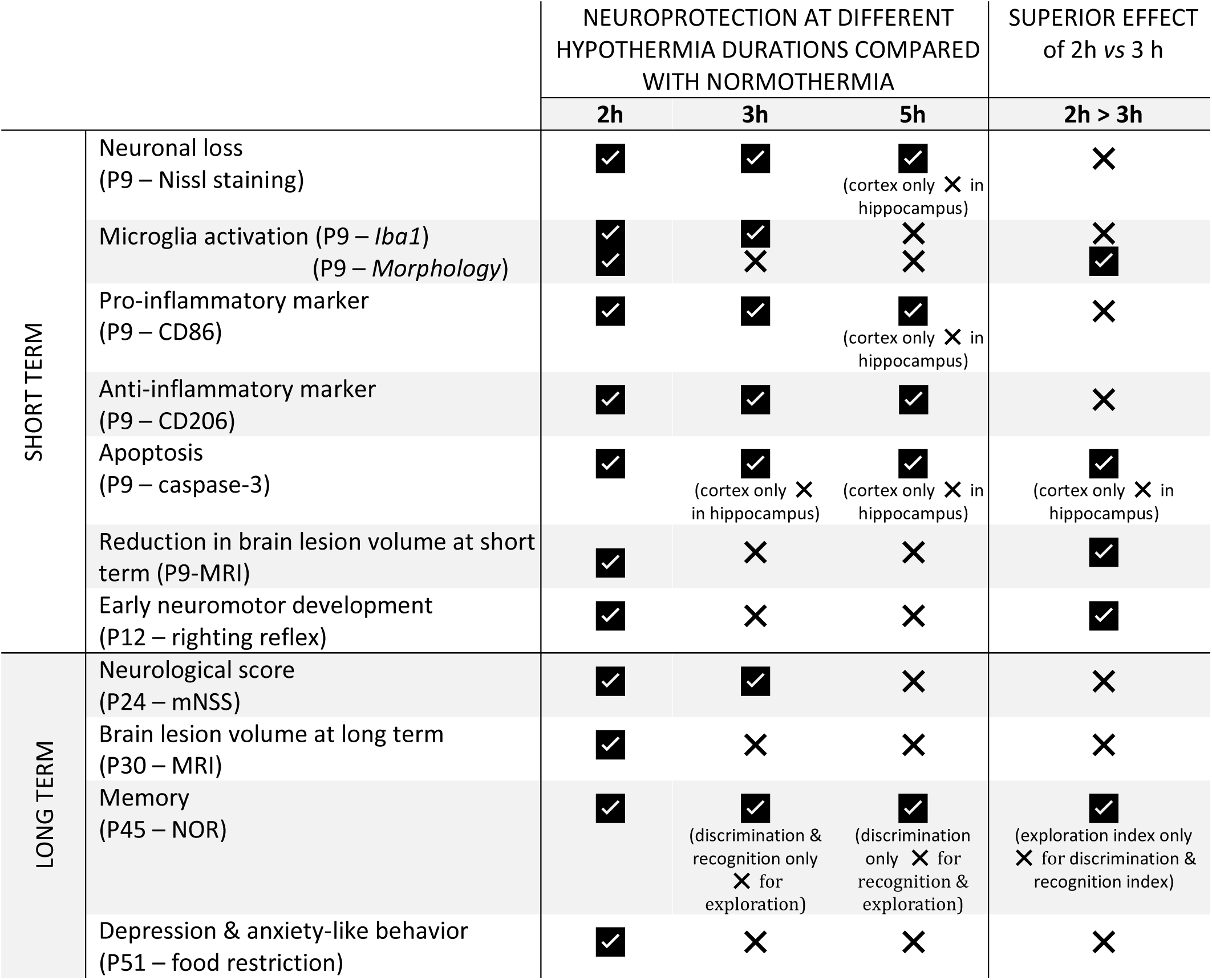
Summary of neuroprotective effects of different hypothermia durations compared with normothermia (HIN group) and relative superiority of 2 h versus 3 h hypothermia. Green symbol indicates a statistically significant difference between the hypothermia-treated group and the normothermia (HIN) group, or a statistically significant difference between 2- and 3-hour hypothermia. Red symbol indicates no statistically significant difference between hypothermia-treated groups and the normothermia (HIN) group, or between 2- and 3-hour hypothermia.

|  |  | NEUROPROTECTION AT DIFFERENT HYPOTHERMIA DURATIONS COMPARED WITH NORMOTHERMIA |  |  | SUPERIOR EFFECT of 2h vs 3 h |
| --- | --- | --- | --- | --- | --- |
|  |  | 2h | 3h | 5h | 2h > 3h |
| SHORT TERM | Neuronal loss (P9 – Nissl staining) | ✓ | ✓ | ✓<br>(cortex only ✗ in hippocampus) | ✗ |
|  | Microglia activation (P9 – *Iba1*)<br>(P9 – *Morphology*) | ✓<br>✓ | ✓<br>✗ | ✗<br>✗ | ✗<br>✓ |
|  | Pro-inflammatory marker (P9 – CD86) | ✓ | ✓ | ✓<br>(cortex only ✗ in hippocampus) | ✗ |
|  | Anti-inflammatory marker (P9 – CD206) | ✓ | ✓ | ✓ | ✗ |
|  | Apoptosis (P9 – caspase-3) | ✓ | ✓<br>(cortex only ✗ in hippocampus) | ✓<br>(cortex only ✗ in hippocampus) | ✓<br>(cortex only ✗ in hippocampus) |
|  | Reduction in brain lesion volume at short term (P9-MRI) | ✓ | ✗ | ✗ | ✓ |
|  | Early neuromotor development (P12 – righting reflex) | ✓ | ✗ | ✗ | ✓ |
| LONG TERM | Neurological score (P24 – mNSS) | ✓ | ✓ | ✗ | ✗ |
|  | Brain lesion volume at long term (P30 – MRI) | ✓ | ✗ | ✗ | ✗ |
|  | Memory (P45 – NOR) | ✓ | ✓<br>(discrimination & recognition only ✗ for exploration) | ✓<br>(discrimination only ✗ for recognition & exploration) | ✓<br>(exploration index only ✗ for discrimination & recognition index) |
|  | Depression & anxiety-like behavior (P51 – food restriction) | ✓ | ✗ | ✗ | ✗ |

## DISCUSSION

Controlled moderate hypothermia is currently the only established intervention for neonatal hypoxic–ischemic injury (NHI), yet nearly 50% of treated infants still develop severe neurological disabilities or die ^46^. This underscores the critical need for preclinical research aimed at identifying novel therapeutic strategies. Importantly, such investigations must benchmark emerging therapies against the established effects of hypothermia to determine whether they offer additional neuroprotective benefit. The objective of this study was therefore not to challenge or modify the clinically established hypothermia protocols, but rather to define a standardized and maximally neuroprotective reference protocol in a preclinical setting. Establishing such a reference ensures that any novel neuroprotective treatment can be reliably evaluated for additional benefit, increasing the likelihood that promising interventions can be translated to the clinic and potentially complement therapeutic hypothermia, which remains the standard of care.

To achieve this, the well-established Rice–Vannucci model was used in postnatal day 7 (P7) rats, which induces a moderate to severe injury ^23,32,47,48^. Rectal temperatures were set at 36.0 ± 0.5 °C for normothermia and 32.0 ± 0.5 °C for hypothermia. The hypothermia temperature of 32 °C was chosen based on existing literature ^23,32,47,48^ and to remain below the normothermia range observed in P7 rat pups bred and housed under modern laboratory conditions (between 33.5°C and 36.0°C) ^34^. The 2°C to 6°C difference below normothermia is considered optimal for therapeutic hypothermia, positioning the study’s temperature differential within this range ^49^. The normothermic temperature value was selected based on measurements of P7 nesting pups, which exhibited a lower baseline temperature compared to the commonly used 37°C in preclinical models ^23,50–52^. This aligns with the findings of Wood *et al*. ^34^, who reported that the median rectal temperature of healthy nesting pups varied from 35.4 °C to 36.1 °C, with temperature variability decreasing between P7 and P14. This is also consistent with our own observations, in which the rectal temperatures of nesting pups with their dams ranged from a median of 35.6°C to 35.9°C. In the normothermic group, rat pup rectal temperature was maintained at 35.8 ± 0.3°C over a 2-hour period, after which they were returned to their dams. Given the considerable variation in the duration of hypothermia reported in preclinical studies, this study aimed to evaluate the effects of some of the most commonly used durations under consistent environmental conditions and within the same hypoxia-ischemia model. Our objective was to define a practical and reproducible protocol suitable for preclinical testing, allowing reliable comparisons with other experimental neuroprotective interventions, in order to ensure that any observed benefit from the additional treatment can be confidently attributed to its therapeutic effect, rather than variability in the therapeutic hypothermia protocol.

A major strength of this study is the use of non-invasive MRI to monitor lesion volumes longitudinally in the same animals, together with behavioral tests, and immunohistochemical analyses. This combined approach let us thoroughly assess how different hypothermia durations affect brain damage, motor and cognitive function, and cellular and molecular outcomes, from neonatal pups all the way to young adults. Short-term effects were assessed using MRI at P7 and P9, behavioral tests at P8, P10, and P12, and histological and immunohistochemical analyses at P9, whereas long-term effects were evaluated by MRI at P30 and behavioral testing up to P51. This longitudinal approach adheres to the STAIR consortium’s recommendations, particularly point 4 ^41^, highlighting the importance of such studies, as early neuropathological markers do not consistently predict long-term outcomes ^53,54^. In this study, we demonstrated that controlled and mild hypothermia lasting 2 hours, maintaining a rectal temperature of 32°C, was sufficient to provide neuroprotection following a neonatal hypoxic-ischemic injury on all the measured parameters (Table 2). Interestingly, extending the duration of hypothermia under identical HI conditions did not lead to additional benefits. On the contrary, the 5h-therapeutic hypothermia protocol was associated with less favorable outcomes overall, despite conferring partial long-term neuroprotection for specific parameters, such as memory performance, compared with the HIN group. Prolonged hypothermia may blunt endogenous recovery processes, increase stress during extended exposure and alter inflammatory responses during rewarming, which together could attenuate neuroprotection. Compared with 3 h of hypothermia, 2 h of therapeutic hypothermia resulted in superior outcomes, especially at short-term, as evidenced by a greater reduction in brain lesion volume within the first 48 h and improved righting reflex performance in pups. Similar findings were observed in a P7 Sprague Dawley model of NHI treated with 2 hours of hypothermia at a lower set temperature of 30.0 ± 0.5°C ^30^. In this latter study, early reflexes were assessed using the geotaxis and cliff aversion tests during the week following the hypoxic-ischemic event. The authors concluded that therapeutic hypothermia could mitigate the detrimental effects of the injury on the reflexes of neonatal rats. In our study, analysis of long-term outcomes showed that both 2 h and 3 h of therapeutic hypothermia were associated with improved outcomes, supporting neuroprotection of motor and cognitive functions in rats. However, the shortest time period of hypothermia led to lower brain lesion volumes at P30 as well as less depression or anxiety-like behavior at P51. Taken together, these findings indicate that 2 h of therapeutic hypothermia is sufficient to confer neuroprotection in both the short and long term, in agreement with previous reports ^55^.

Although some preclinical studies in P7 rats have reported sex-related differences following hypoxic-ischemic injury ^35,38^, we did not observe any significant sex differences in lesion volumes or behavioral test performance across the different experimental groups. Similar findings showing no sex-specific differences have been reported in other studies using the Rice-Vannucci model in neonatal rats ^48,56^. Furthermore, a study in piglets also found no evidence of sex-specific differences in brain lesions following NHI ^57^. However, clinical research suggests that male infants may be more vulnerable to brain injury and exhibit more pronounced long-term neurological deficits than female infants after NHI ^58,59^. Other evidence from the literature highlights a sexual dimorphism in the expression of some proteins linked to the cell death pathway, for example ^60–62^. Notably, one study reported a more pronounced reduction in caspase-3 activation in females compared to males following a unilateral focal ischemia-reperfusion injury in P7 rats ^60,61^. Given the temperature sensitivity of the caspase-dependent apoptotic pathway, females may derive greater neuroprotective benefit from hypothermic treatment. Therefore, although our data do not reveal sex-specific differences, this absence of effect might be model- or sample-size dependent, and highlights the need for further research into sex-dependent responses to hypothermia, especially in relation to apoptotic pathways. Indeed, one limitation of our study is the small number of animals used for immunohistochemical and histological analyses, such as the aforementioned potential sex dimorphism in caspase-3 expression. However, this limitation reflects our methodological choice to prioritize a longitudinal *in vivo* approach, combining non-invasive MRI with behavioral assessments. This strategy allows for tracking brain–behavior dynamics over time, acknowledging that MRI alone is not always predictive of functional outcomes, particularly in the context of high neuroplasticity in neonatal models ^53,54^.

Consistent with previous reports ^60,61^, caspase-3 activation was observed in all groups following NHI. All hypothermia treatments significantly reduced caspase-3 activation in the cortex, with the HIH2 group showing a greater reduction than HIH5 in both cortex and hippocampus. In the hippocampus, only 2 hours of hypothermia significantly decreased apoptosis compared to the other HI groups, in line with improved performance in the novel object recognition test, reflecting hippocampus-dependent memory. Consistent with these findings, Nissl staining revealed that cortical neuronal loss was prevented in all hypothermia-treated groups, with the HIH2 group exhibiting greater neuronal preservation than HIH5. In the CA1 region of the hippocampus, neuronal preservation was more pronounced following 2 h and 3 h hypothermia.

The suppression of microglial pro-inflammatory activity is recognized as a beneficial marker ^63^ and a critical mechanism by which therapeutic hypothermia exerts its neuroprotective effects ^64^. In order to strengthen the analysis of the different time duration of hypothermia, we decided to perform immunohistochemistry in two brain structures, the cortex and the CA1 area of the hippocampus, as these cerebral regions are the most affected in the NHI animal model ^35,38^. In this study, two hours of hypothermia led to a marked decrease in Iba1 fluorescence intensity in both the cortex and the CA1 region, indicating reduced microglial activation. This reduction was less significant after three hours of hypothermia and completely lost after five hours of therapeutic cooling. In addition, in the cortex and the CA1 region of the hippocampus, NHI led to a marked reduction in microglial arborization complexity, as evidenced by decreases in the number of branches, junctions, and branching points, indicative of a transition toward an activated, amoeboid microglial phenotype ^65^. Remarkably, a 2-hour hypothermia treatment prevented these structural alterations, maintaining a highly ramified morphology, that was not the case for longer hypothermia durations. These findings are consistent with previous studies showing that resting microglia exhibit extensive branching, whereas activated microglia are characterized by shortened branches and enlarged cell bodies, aligning with the microglial morphology observed following NHI in our study ^66,67^. Furthermore, 2-, 3-, and 5-hour hypothermia treatments modulated the microglial phenotype, leading to decreased expression of the pro-inflammatory marker CD86 in the cortex. However, in the CA1 region of the hippocampus, the decrease in CD86 expression was measured only after shorter hypothermia durations, 2 and 3 hours. In addition, all hypothermia-treated groups showed increased expression of the anti-inflammatory marker CD206, indicating a shift toward a more anti-inflammatory microglial profile. A previous study reported that hypothermia not only reduced microglial activation 48 hours after HI injury ^55^, but also decreased the expression of the pro-inflammatory marker CD86, reaching levels comparable to those of healthy sham mice. It was also shown that hypothermia further enhanced the expression of the anti-inflammatory marker CD206 at both 3 and 7 days after neonatal HI ^68^.

Taken together, these findings demonstrate that prolonged durations of hypothermia do not confer additional neuroprotective benefit over shorter protocols, as a 2-hour hypothermia treatment is sufficient to attenuate microglial activation, promote an anti-inflammatory phenotype, and reduce apoptotic and neuronal damage following NHI. These observations support the existence of an optimal therapeutic window in which moderate hypothermia preserves microglial homeostasis and neuronal integrity after hypoxic–ischemic injury. Prolonged cooling may, however, engage physiological or cellular responses that counteract neuroprotection. One plausible mechanism is increased stress associated with extended hypothermia exposure in immature rodents, as prolonged cooling has been reported to induce agitation, shivering, or restraint-related stress responses that may interfere with the beneficial effects of hypothermia ^56,69^. In addition, stress following hypoxic–ischemic injury has been shown to induce morphological and phenotypic alterations in microglia in the rodent brain ^70^. Consistent with this interpretation, modulation of inflammatory and apoptotic markers was less pronounced in the HIH5 group than in the HIH2 and HIH3 groups. Further mechanistic studies, including assessment of stress-related biomarkers, controlled rewarming paradigms, and temporal profiling of inflammatory and apoptotic signaling, will be required to delineate these interactions.

This study also compared the neuroprotective effects of initiating therapeutic hypothermia either immediately after exiting the hypoxic chamber (HIH2Im) or 1 ½ hour later (HIH2). In the short term, immediate hypothermia reduced lesion volumes more than delayed treatment, although this difference diminished over time, with comparable lesion sizes at P30. Motor and reflex recovery were similar between groups, but early hypothermia improved performance in the novel object recognition test, reflecting enhanced long-term memory. These findings regarding the long-term memory neuroprotection align with previous research by Sabir *et al*., who reported a linear decrease in neuroprotection with increasing delay in the initiation of hypothermia in a P7 hypoxic-ischemic pup model ^37^. Although initiating therapeutic hypothermia earlier yields a slightly stronger neuroprotective effect in our pre-clinical study, this conclusion is based primarily on early MRI and the novel object recognition test, as no histological analyses were performed in HIH2Im and no differences were observed neither in the other behavioral tests, nor with MRI at P30. However, this tendency is consistent with clinical trials indicating that newborns experiencing hypoxic-ischemic insults had better outcomes when cooling began within 3 hours post-event, compared to those whose treatment was initiated between 3 and 6 hours post-hypoxia-ischemia ^71,49^.

Overall, our data suggest that 2 hours of hypothermia is the most effective duration in our rat model. To relate this to clinical protocols, the differences between rats and humans must be considered. While rat physiology is often regarded as closer to humans than mice, their biological timescales differ markedly. A “rat hour” or “rat day” does not correspond directly to a human hour or day, as rats live roughly 27 times faster than humans ^72^. This implies that 72 hours of hypothermia in human neonates roughly corresponds to 2–3 hours in rats. Nevertheless, such extrapolations should be interpreted cautiously due to developmental and physiological differences.

## CONCLUSION

Although hypothermia is the standard clinical care for newborns with moderate-to-severe hypoxia-ischemia in high-income countries, this treatment has limitations in eligible neonates, as 44-53% do not respond favorably ^15^. Moreover, advanced technologies for delivering hypothermia and comprehensive monitoring of affected infants often remain unavailable in many hospitals in low- and middle-income countries ^73^. For instance, while many pediatricians in South Africa recognize the effectiveness of hypothermia in reducing neurological deficits in NHI, fewer than half are able to offer it as a treatment option ^74^. This highlights the urgent need to develop new treatment options, which must be compared to hypothermia.

Refining preclinical models in the pursuit of new neuroprotection strategies for neonatal hypoxia-ischemia is therefore critical. It allows researchers (i) to determine the optimal conditions for combining hypothermia with other neuroprotective approaches, balancing their benefits and minimizing potential side effects, and (ii) to directly compare hypothermia as a stand-alone treatment with other therapeutic strategies. This latter point is particularly relevant in preclinical research. However, it is important to note that hypothermia efficacy likely depends on factors such as injury severity, timing, and biological differences between species. While our findings indicate that 2 hours of hypothermia at 32°C confers the best neuroprotective outcomes under the conditions tested in Wistar rat pups, further studies might be required to refine the optimal protocol in other strains or species, while minimizing side effects related to animal welfare, thereby addressing the third principle of the 3Rs (Replacement, Reduction, Refinement).

## DATA AVAILIBILITY

The datasets generated during and/or analyzed during the current study are available from the corresponding author upon request.

## ACKNOWLEDGEMENTS

The present study was conducted in the context of the framework of the University of Bordeaux’s IdEx‘Investments for the Future’ program/RRI“IMPACT.” This work was supported by the Fédération pour la Recherche sur le Cerveau (FRC) and by the Nouvelle Aquitaine region. This work received also financial support from the French Agence Nationale de la Recherche (ANR) grant, BrainFuel reference (grant number ANR-21-CE44-0023 to A.K.B.S. and L.P.) and the Spark program.

## STATEMENTS AND DECLARATIONS

Authors have no conflict of interest to declare.

## AUTHOR CONTRIBUTION

The last two authors contributed to the study conception and design. Experiments, data collection and analyses were performed by Ifrah Omar, Pierre Goudeneche, Léonie Dayraut, Hélène Roumes, and Anne-Karine Bouzier-Sore. The first draft of the manuscript was written by Hélène Roumes, Ifrah Omar, and Anne-Karine Bouzier-Sore. All authors commented on previous versions of the manuscript. All authors read and approved the final manuscript.

## Supplementary Figure Legends

**Supplementary Fig. S1:**
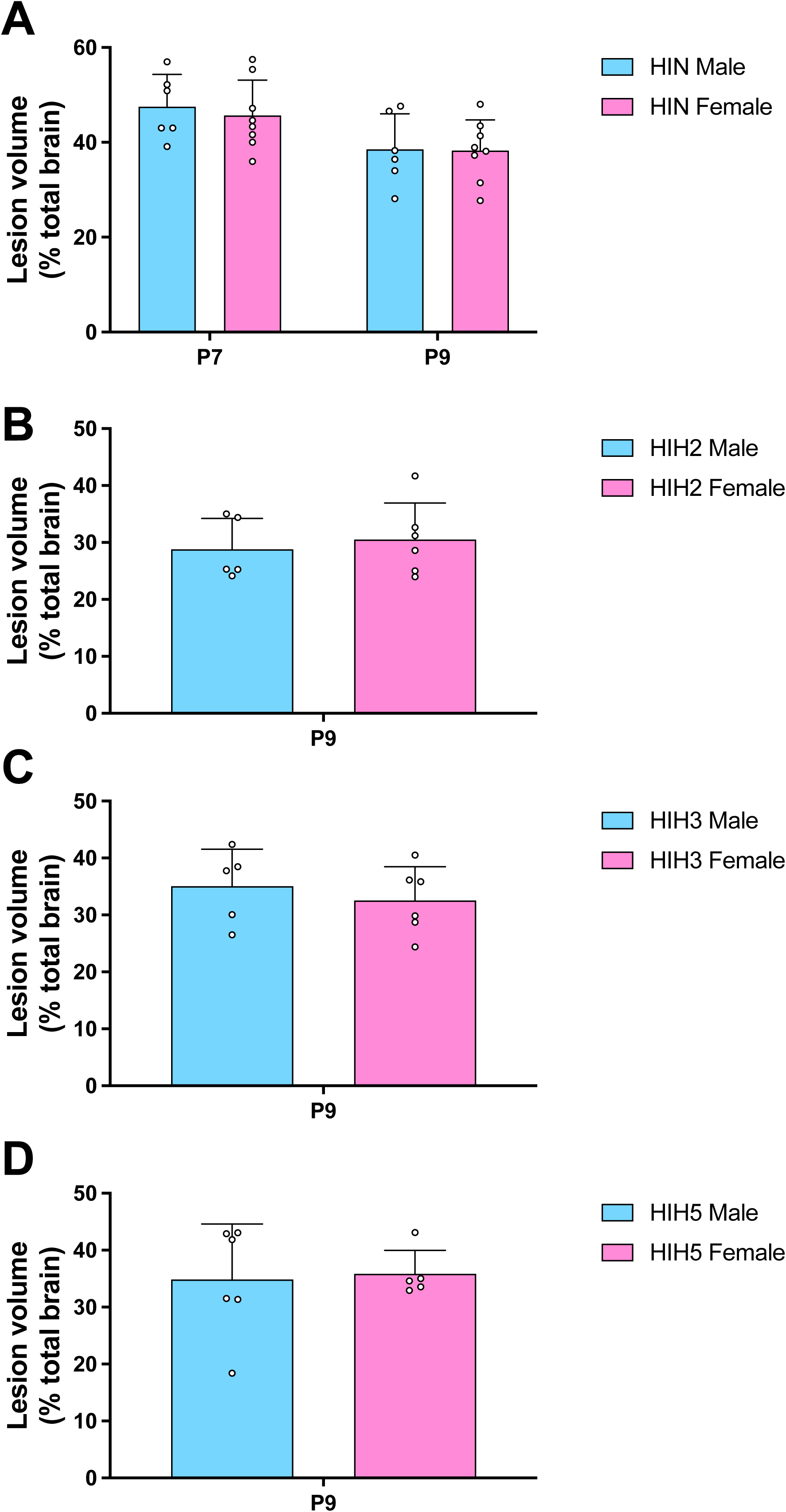
Sex-comparison of brain lesion volumes at short term. (A) HIC groups, at P7 and P9. (B) HIH2 group at P9. (C) HIH3 group at P9. (D) HIH5 group at P9.

**Supplementary Fig. S2.**
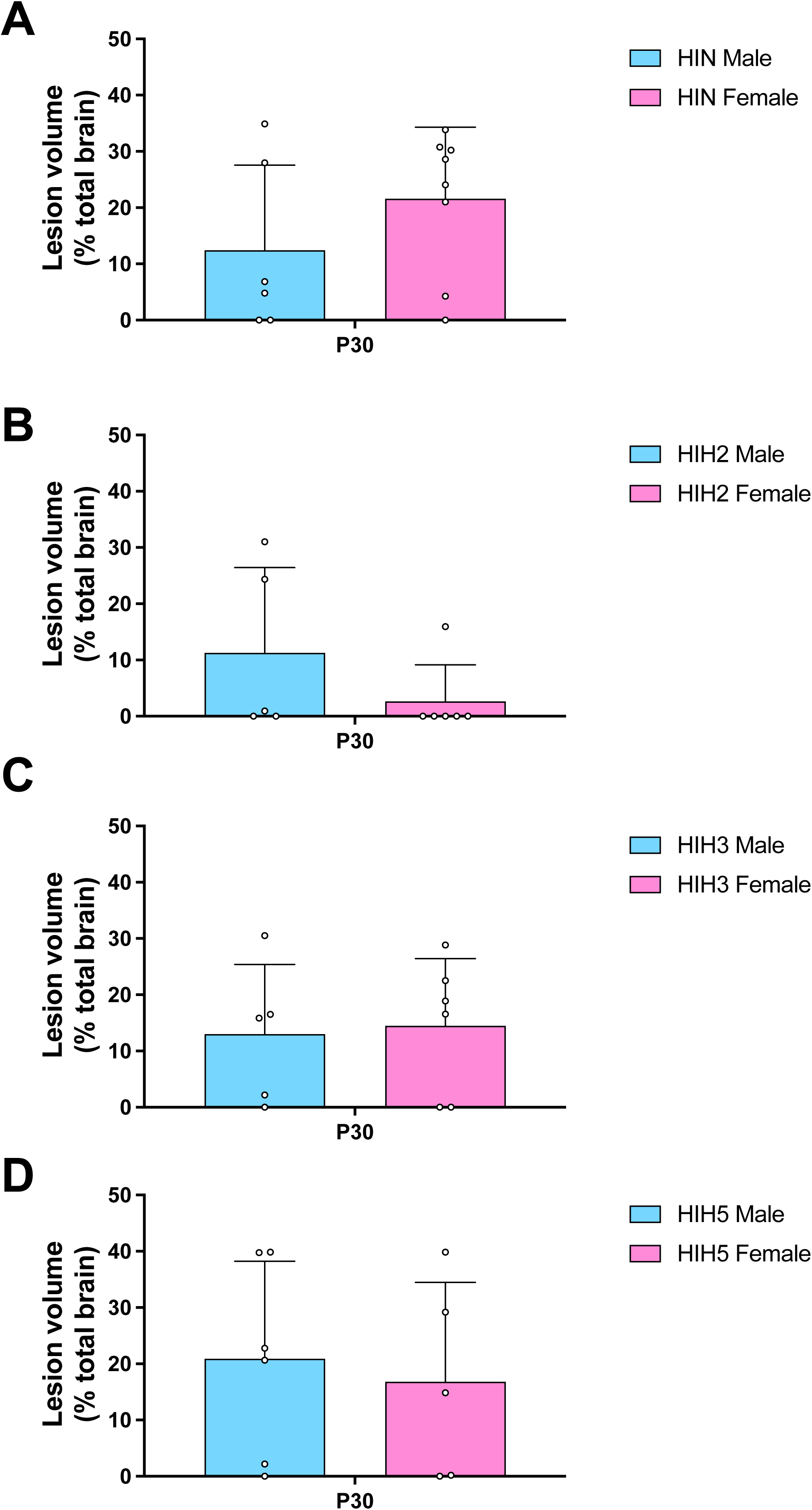
Sex-comparison of brain lesion volumes at long term. (A) HIC groups, at P30. (B) HIH2 group at P30. (C) HIH3 group at P30. (D) HIH5 group at P30.

**Supplementary Fig. S3:**
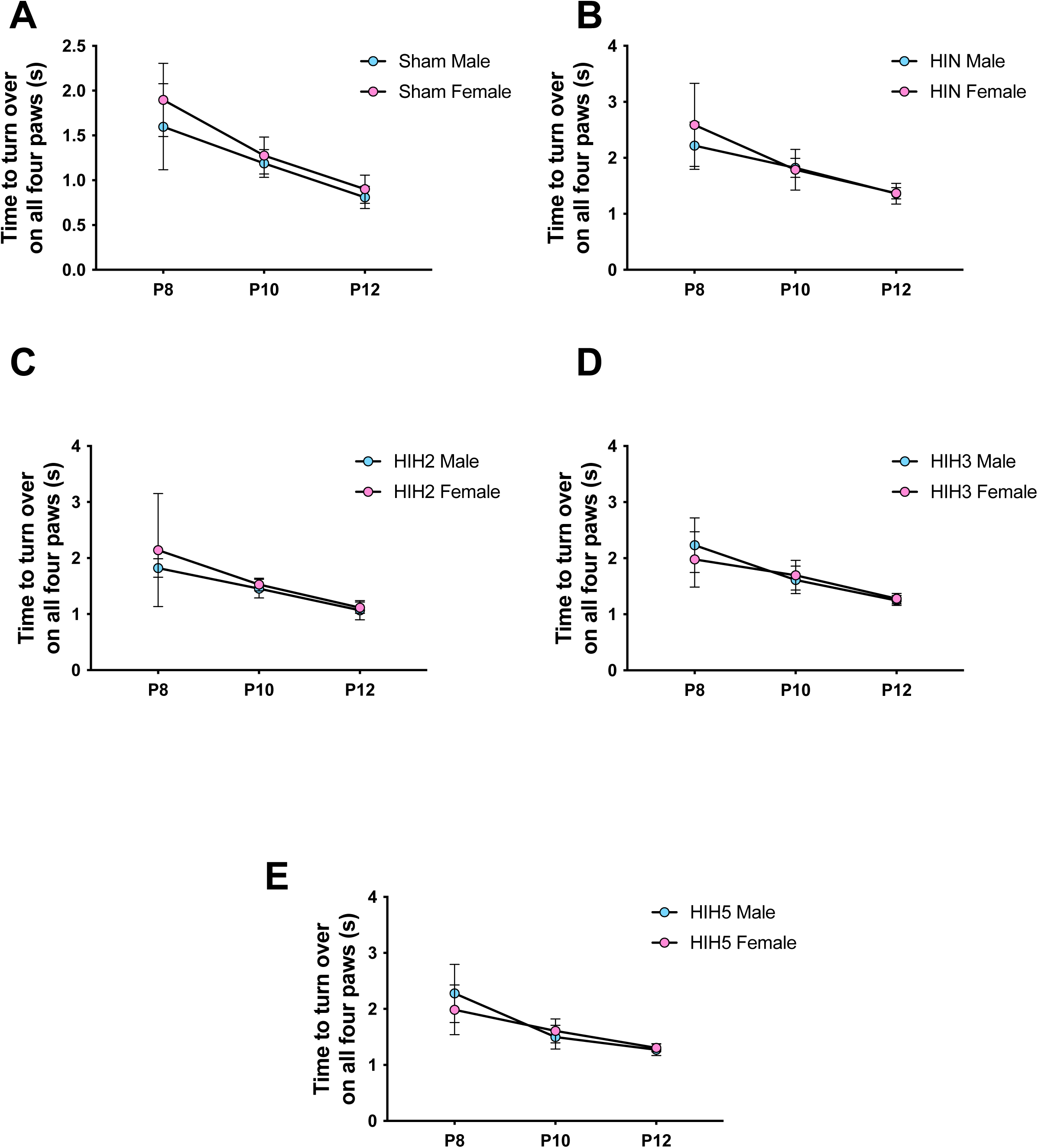
Sex-comparison for the early righting reflex test. (A) Sham groups (B) HIN group. (C) HIH2 group. (D) HIH3 group. (E) HIH5 group.

**Supplementary Fig. S4:**
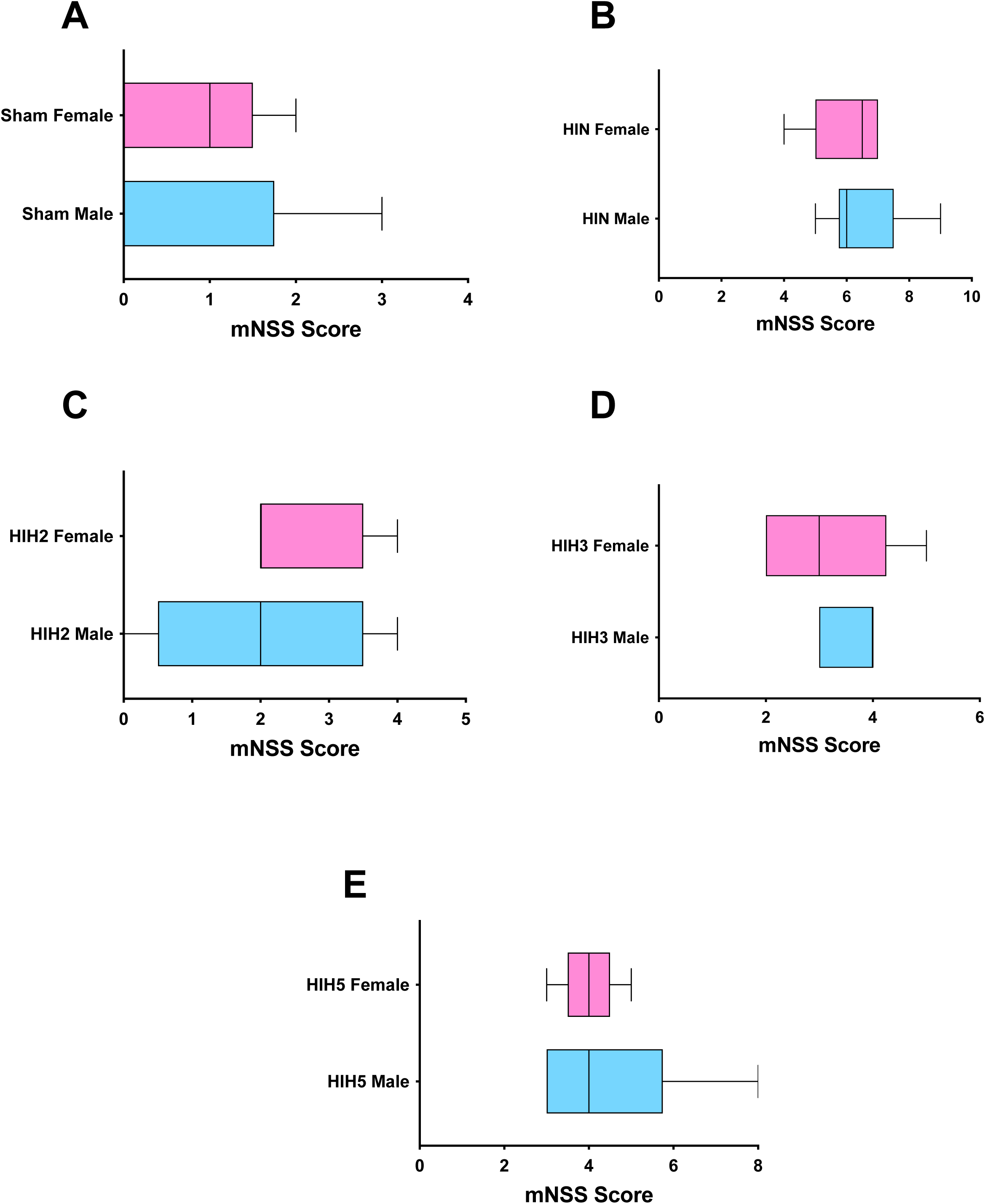
Sex-comparison for the modified neurological severity scores. (A) Sham group. (B) HIN group. (C) HIH2 group. (D) HIH3 group. (E) HIH5 group.

**Supplementary Fig. S5:**
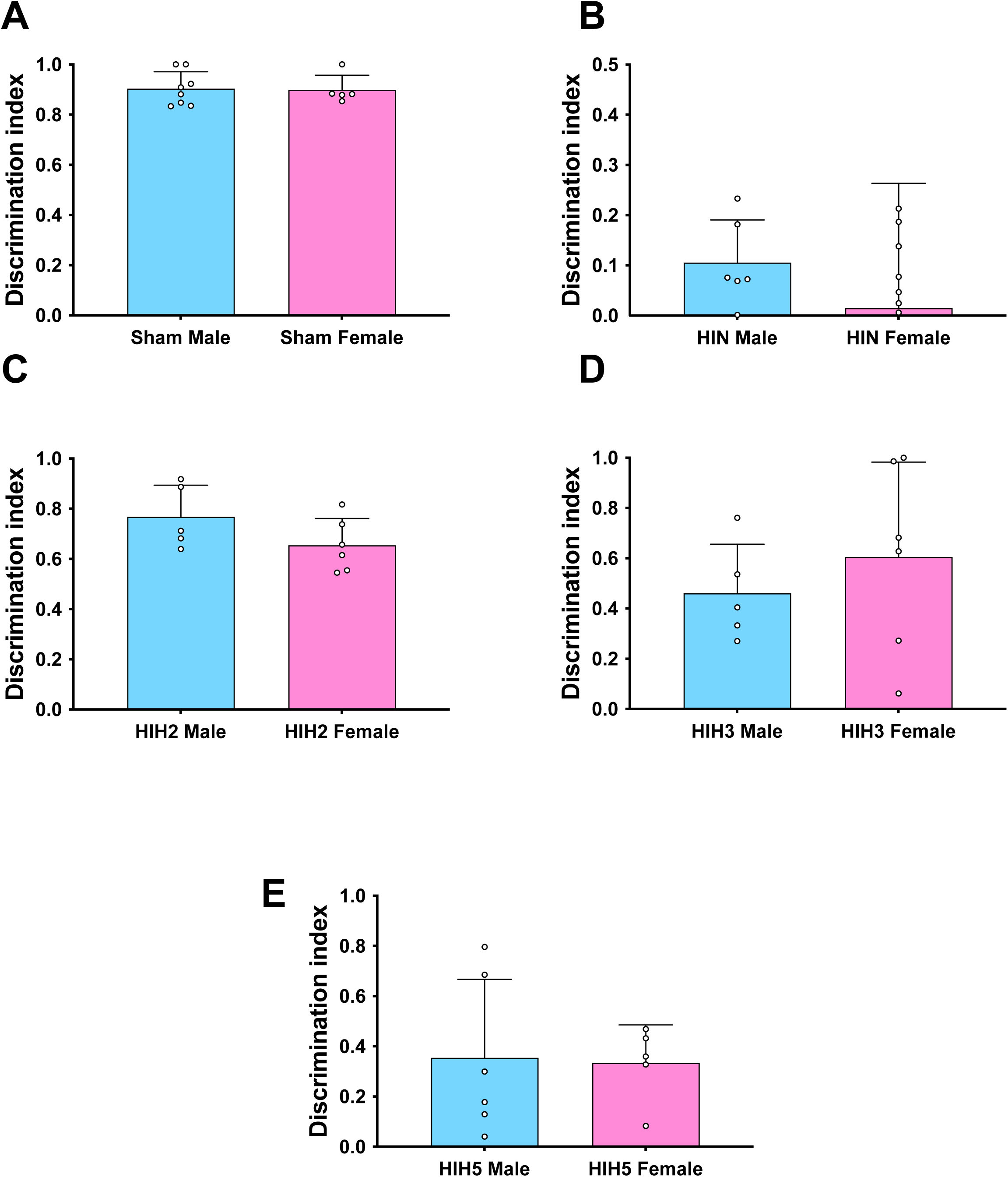
Sex-comparison for the novel object recognition test performed at P45. (A) Sham group. (B) HIN group. (C) HIH2 group. (D) HIH3 group. (E) HIH5 group.

**Supplementary Fig. S6:**
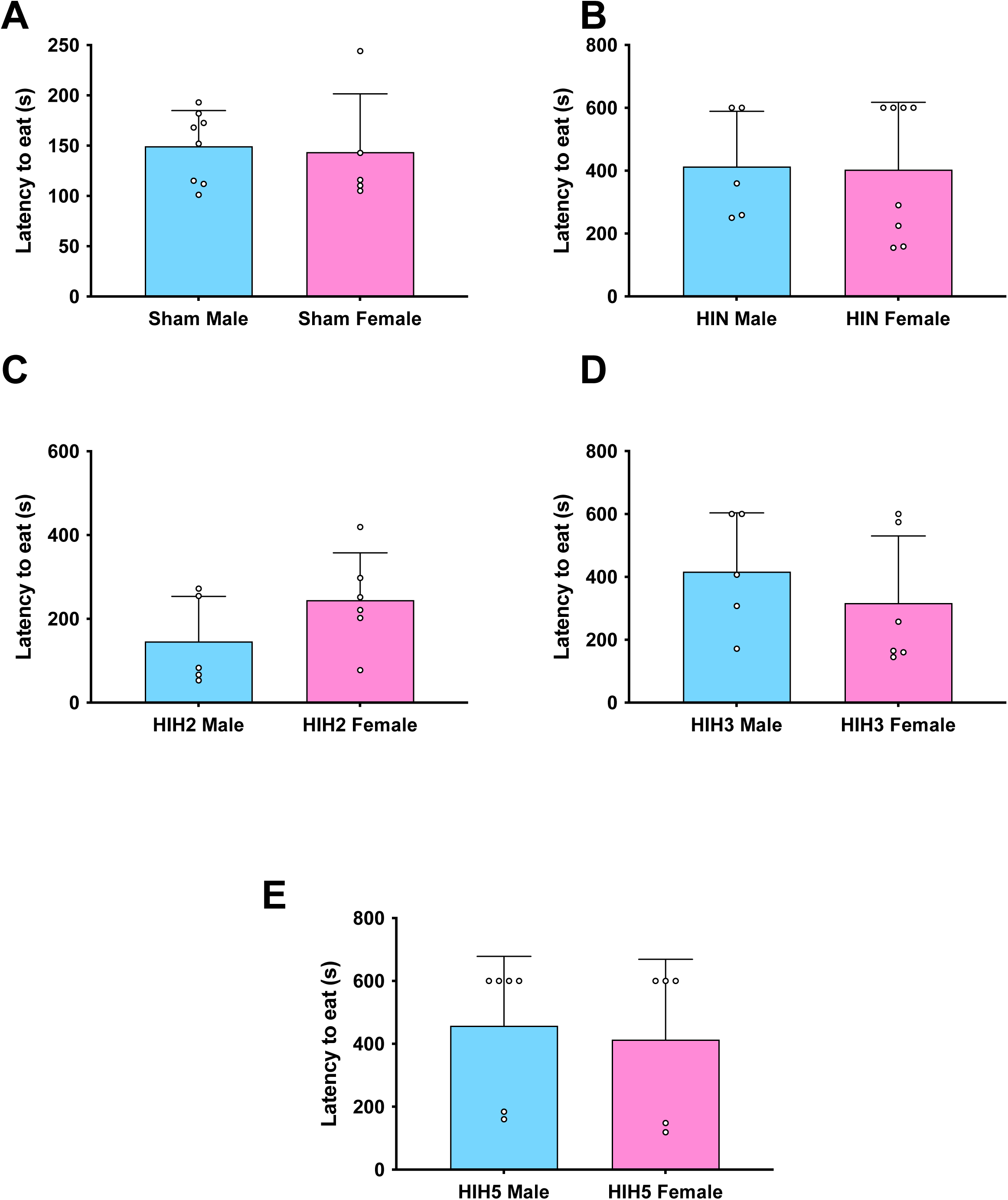
Sex-comparison for the food-restriction test performed at P51. (A) Sham group. (B) HIN group. (C) HIH2 group. (D) HIH3 group. (E) HIH5 group.

**Supplementary Fig. S7:**
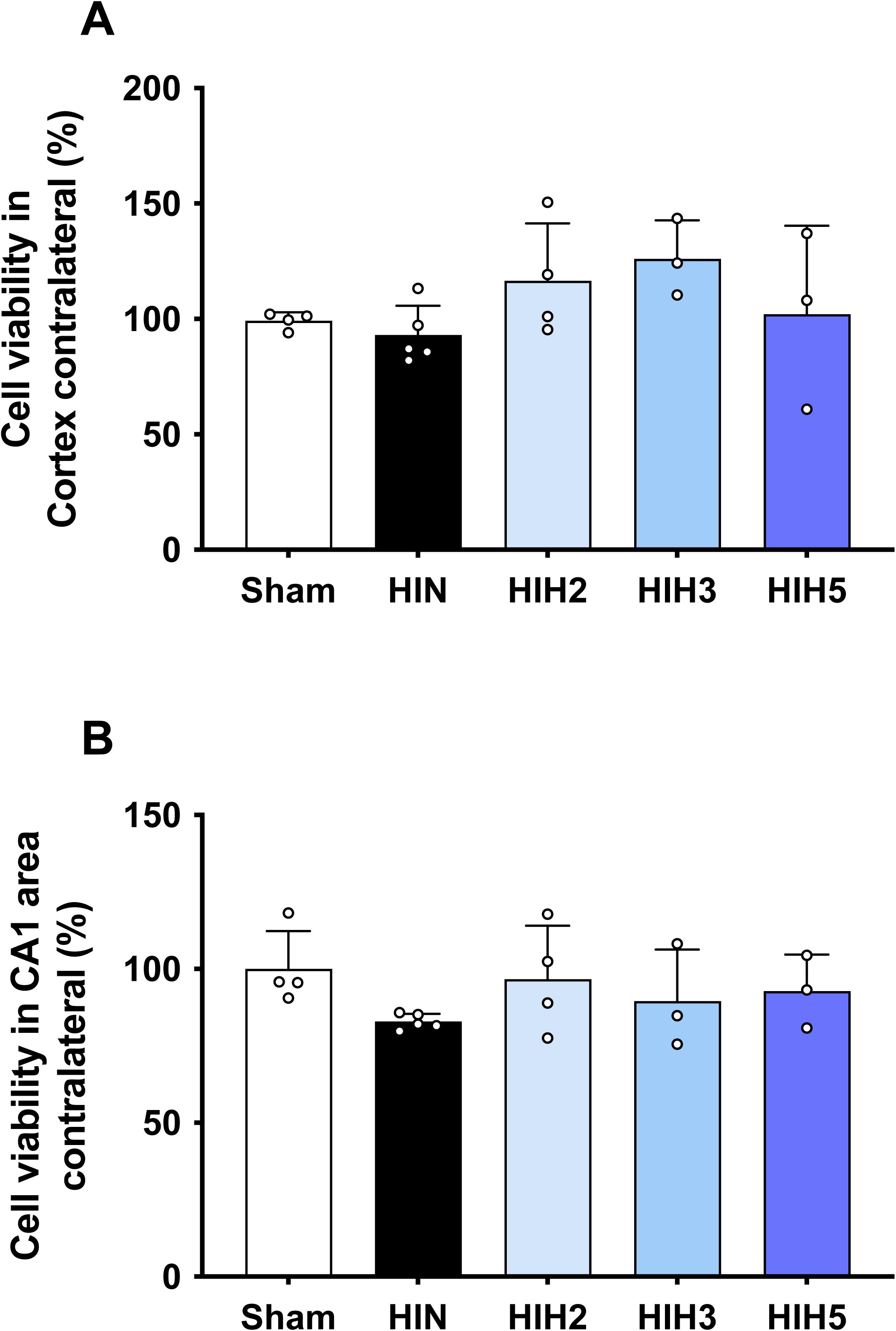
Histological quantification of cell viability using Cresyl violet staining. Cell viability (% related to Sham) in contralateral brain cortex (A) and in contralateral hippocampus (B).

**Supplementary Fig. S8:**
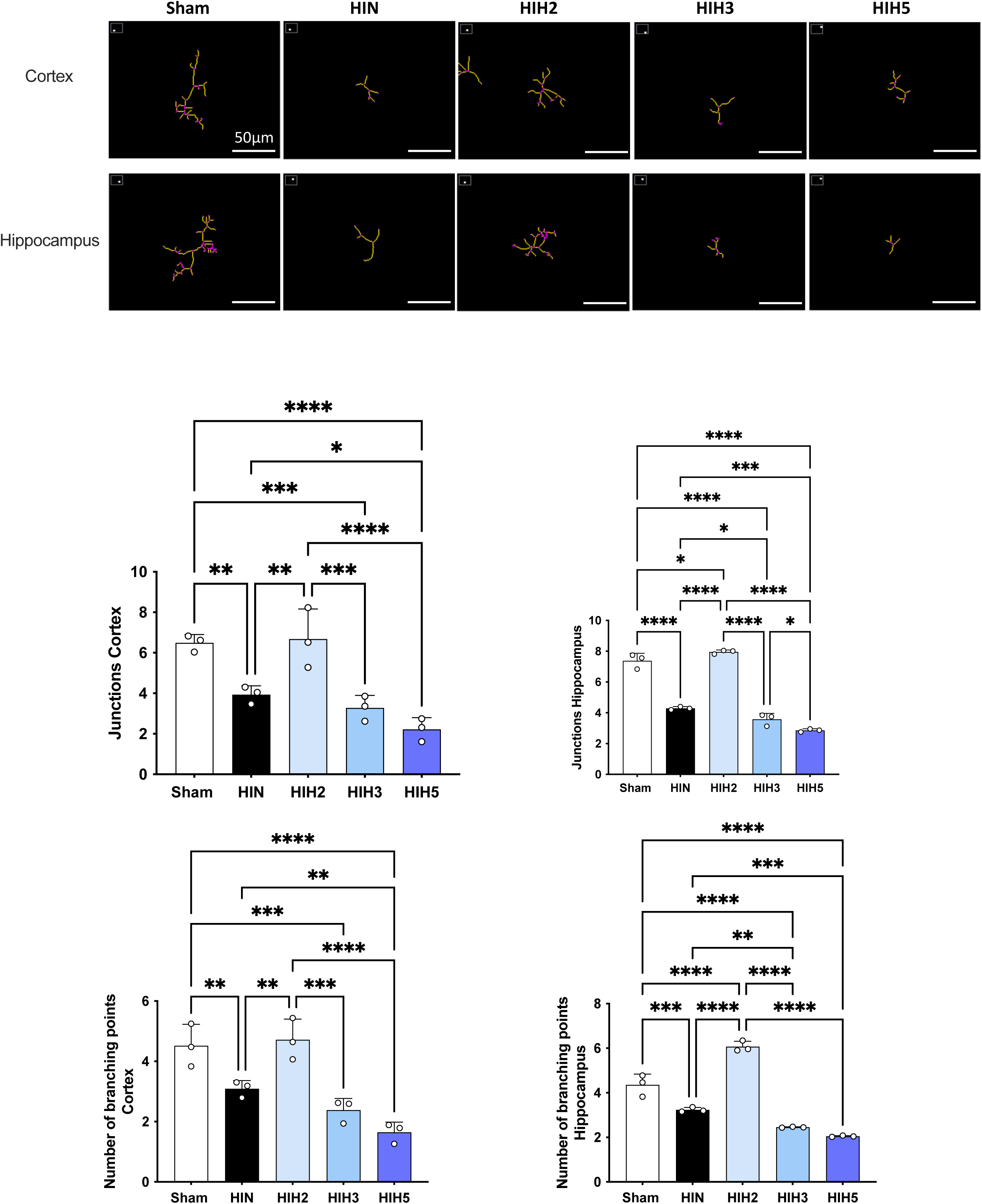
Impact of a 2h-hypothermia treatment after NHI on microglia morphology. Top: Representative images of skeletonized microglia in the cortex and hippocampus, with branch junctions highlighted in violet. Bottom: Quantification of microglial morphology showing the mean number of branch junctions and quadruple points in the cortex and hippocampus across the 5 experimental groups.

